# Pneumococcal competence, a populational health sensor driving multilevel heterogeneity in response to antibiotics

**DOI:** 10.1101/2023.11.17.567514

**Authors:** Marc Prudhomme, Calum H. G. Johnston, Anne-Lise Soulet, David De Lemos, Nathalie Campo, Patrice Polard

## Abstract

Competence for natural transformation is a central driver of genetic diversity in bacteria. In the human pathogen *Streptococcus pneumoniae*, competence exhibits a populational character mediated by the stress-induced ComABCDE quorum-sensing (QS) system. Here, we explore how this cell-to-cell communication mechanism proceeds and the functional properties acquired by competent cells grown under lethal stress. We show that populational competence development depends on self-induced cells stochastically emerging in response to stresses, including antibiotics. Competence is demonstrated to propagate through the population from a low threshold density of self-induced cells, defining a biphasic Self-Induction and Propagation (SI&P) QS mechanism. We also reveal that a competent population displays either increased sensitivity or improved tolerance to lethal doses of antibiotics, dependent in the latter case on the competence-induced ComM division inhibitor. Remarkably, these surviving competent cells also display an altered transformation potential. Thus, the unveiled SI&P QS mechanism shapes pneumococcal competence as a health sensor of the clonal population, promoting a bet-hedging strategy that both responds to and drives cells towards heterogeneity.

## Introduction

In all kingdoms of life, collective behaviour provides novel properties to a population unattainable at the individual level. Effective communication between individuals is key to achieving these features. In bacteria, such communication systems are crucial for survival, enabling an adaptive response to the selective pressure exerted by the niche on the population ^1,2^. A widespread mode of bacterial cell-to-cell communication is quorum-sensing (QS), which relies on small exported molecules called autoinducers (AI). Historically, QS systems were shown to coordinate a cell population once the AI reaches a threshold, inducing gene expression changes and allowing an “all-at-once” populational response to environmental signals ^3^. However, emerging studies on diverse QS systems have highlighted heterogeneity in their mechanisms, promoting differential coordination of the cell population ^4–6^. Unravelling what drives the variability in such QS mechanisms is key to understanding the behaviour of heterogeneously coordinated individuals. Here we addressed these two interlinked questions for the QS system that drives competence for natural transformation in *Streptococcus pneumoniae* (the pneumococcus).

Natural transformation is a key horizontal gene transfer mechanism in prokaryotes, unique in being entirely directed by the recipient cell. It involves the uptake of exogenous DNA followed by its chromosomal integration by homologous recombination, promoting intra- and inter-species genetic exchange ^7,8^. In bacteria, transformation occurs during competence, which is developed and regulated in a species-specific manner ^7^. Pneumococcal competence is transient and is controlled by the ComABCDE QS system, the AI of which is the exported Competence Stimulating Peptide (CSP) ^9^. Remarkably, competence transcriptionally modulates up to 17% of the ∼ 2,000 genes of the pneumococcal genome ^10–12^. Pneumococcal competence provides cells with properties other than transformation, including the ability to kill non-competent relatives, defined as fratricide, which can be mediated by a competence-induced cell-wall hydrolase, CbpD ^13^. Competent cells protect themselves from CbpD via production of the immunity protein ComM ^14^, which also promotes a transient division delay ^15^. In addition, other properties are linked to competence, with the links undefined molecularly but important for the lifestyle of this pathogen in its human host. These include biofilm formation ^16,17^, virulence ^18–22^, increased survival upon transient antibiotic exposure ^23^ and host transmission ^24^.

The pneumococcal competence regulation cycle is well characterized at the single cell level. It can be followed by the exported CSP, expressed from *comC* as a pre-peptide and matured by ComAB during its export. CSP activates the histidine kinase ComD, which in turn phosphorylates its cognate response regulator ComE (Figure 1A) ^25–28^. ComE∼P induces expression of the early competence regulon, which includes the *comABCDE* genes, generating a positive feedback loop ^28^. Two early competence genes encode the alternative sigma factor σ^X^ ^29^ that activates the late competence regulon, including 16 genes required for transformation. Among these is DprA, a conserved late competence protein key for transformation ^30^, which also promotes pneumococcal competence shut-off after ∼30 minutes (Figure 1AB) ^31,32^.

**Figure 1:**
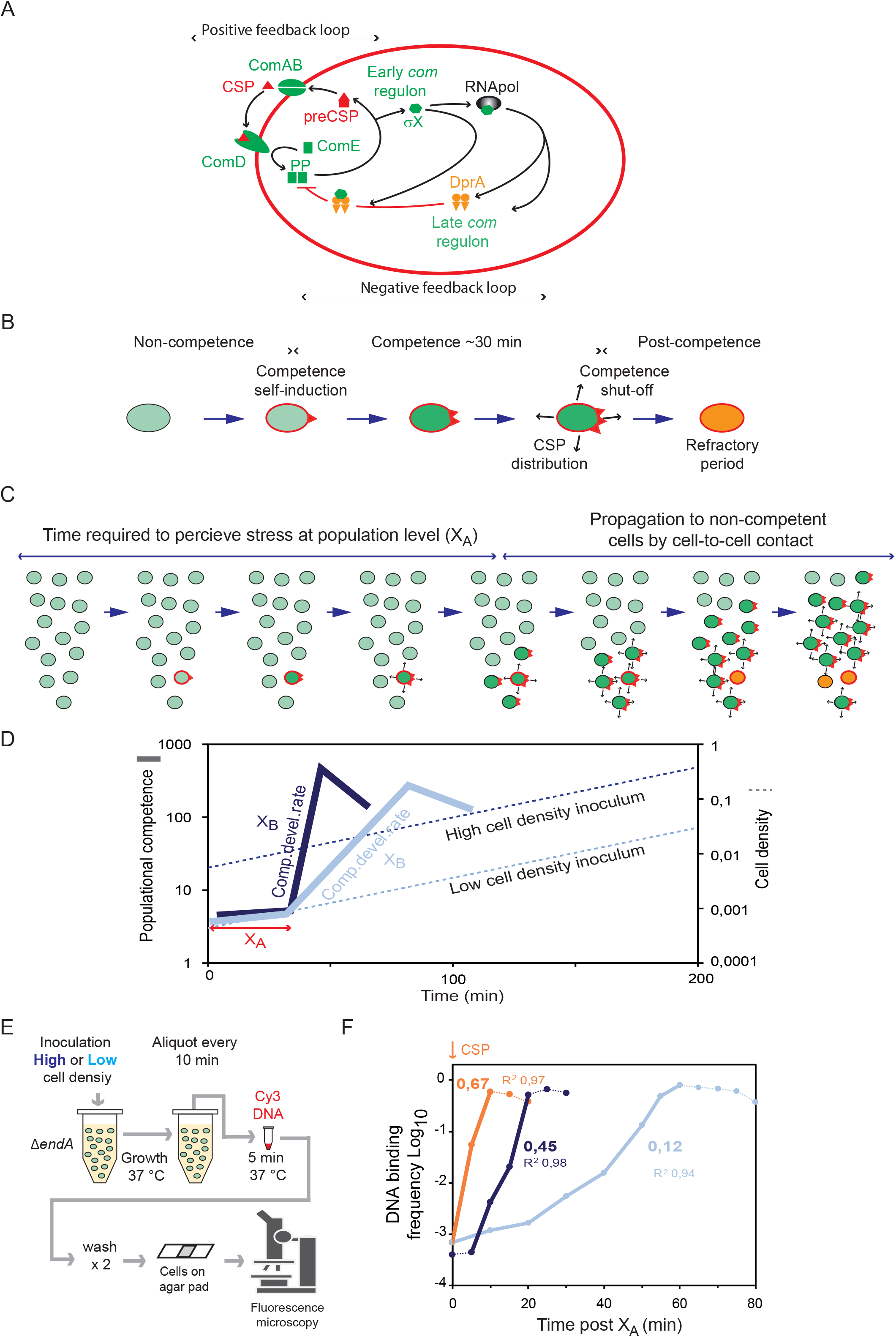
Competence development in the pneumococcus. (A) Model of competence induction at the individual cell level. Pre-CSP is matured and exported by ComAB and interacts with the histidine kinase ComD at the cell pole. ComD phosphorylates its cognate response regulator ComE and ComE∼P dimers induce expression of the early com regulon, including both *comX* genes, which encode the alternative sigma factor σ^X^. σ^X^ interacts with RNA polymerase to induce expression of late com genes, including *dprA*. DprA is then driven to the cell pole by σ^X^ and interacts with ComE∼P to promote competence shut-off. (B) Competence development at the individual cell level. A non-competent cell (black contour, light green fill) senses a stress (red contour) and becomes competent (red contour, dark green fill). This cell becomes a donor of CSP, able to transmit competence to neighbouring cells by cell-to-cell contact (black arrows) and CSP (red triangles) transmission. After ∼30 min, the cell exits competence and enters a post-competence period (red contour, orange fill), during which it is unable to respond to a competence signal (refractory period), before becoming once again able to respond to competence signals. (C) Model of competence development by propagation. In a growing planktonic cell culture in permissive medium, stress and metabolic heterogeneity creates a subpopulation of competent cells able to propagate competence to non-competent neighbouring cells. These neighbouring cells become competent and can in turn transmit the competence signal, leading to exponential propagation throughout the population. After competence, cells enter a post-competence period where they are non-responsive to CSP. Cell identities as in panel B. (D) Model of spontaneous propagation reported by the luciferase transcriptional fusion under the P*_ssbB_* late competence promoter (P*_ssbB_*::*luc*) ^73^. A time (X_A_) is required to produce a cell fraction developing competence in an autocrine mode, which depends on environmental conditions and genotype. The competence development rate (X_B_) is conditioned by the speed of propagation among non-competent cells, which is linked to cell density and the ability to retain CSP. Dotted lines, cell growth; full lines, competence development. (E) Competence propagation visualised by microscopy. Observing binding of fluorescent tDNA to competent cells in different densities of inocula supports the propagation model. TD290 *(endA*^-^) was used to visualize single competent cells ^41^. TD290 was grown in non-permissive C+Y medium and inoculated in permissive C+Y medium at 8 10^-3^ (high cell density, dark blue) or 8 10^-4^ (low cell density, blue) to allow microscopic observation of binding of fluorescent Cy3-DNA by competent cells. (F) The graph reports the frequency of cells binding Cy3-labelled tDNA throughout the culture from the X_A_ time (30 min, Figure S5A), when spontaneous competence induction occurs. This represents a snapshot of the proportion of competent cells in the population at each time point during growth. Light blue curve, low cell density inoculum; dark blue curve, high cell density inoculum; orange curve, low cell density inoculum with CSP added at 0 min (X_A_ time). The rate of increase of cells binding tDNA is reported with the calculated R^2^. Thick lines show data range used for rate calculations. Individual data shown representative of triplicate repeats showing similar results.

Another important characteristic of pneumococcal competence is its induction in exponentially growing cells in response to multiple types of exogenous stress. This key feature was unveiled with planktonic cells grown in an acidic medium, which prevents spontaneous competence development. In these ‘non-permissive’ conditions, addition of exogenous stress can promote competence development ^33–37^. However, how competence is coordinated within a population, either under ‘permissive’ conditions or in response to stress, remains to be defined. In both cases, competence develops over time, unlike upon addition of synthetic CSP where competence is artificially synchronized. The most recent studies on spontaneous competence development, conducted in different strains, reported either a synchronised ^38^ or propagative ^39^ mechanism of populational competence development.

In this study, by combining genetic and single-cell analyses, we revisit the mechanism of competence development. We reveal a general mechanism of competence development irrespective of strain identity and for both spontaneous and stress-mediated competence. In this mechanism, the ComABCDE QS system can stochastically induce competence in response to stress in individual cells of a clonal population, which may lead to propagation through the population. Pneumococcal competence is thus a populational health sensor. We also show that this QS system spreads the clonal population to multilevel heterogeneity, generally increasing tolerance to lethal doses of antibiotics and transformation potential. In all, our results support the view of pneumococcal competence as more than a short-lived program promoting natural transformation, revealing it as a bet-hedging strategy functionally diversifying a clonal population to face multiple types of environmental changes.

## Results

### Spontaneous pneumococcal competence occurs by propagation

The mechanism by which competence develops within a population is still debated, with one study in capsulated pneumococci supporting the historical view of competence as a classical QS system where CSP diffuses and the whole population induces competence in a synchronous manner once a threshold is reached ^38^. In contrast, we showed that competence propagates exponentially through a population by cell-to-cell contact in unencapsulated pneumococci ^39^. This implies the existence of a subpopulation of cells initiating competence, suggesting a bimodal mode of development. We proposed that heterogeneous stress levels in a population stochastically create this subpopulation, which perceives enough stress to self-induce competence linked to CSP retention on the envelope (Figure 1C). This model promotes an exponential wave of competence through the population. We identified two key parameters, a fixed competence induction time (X_A_) associated to the nature of the permissive environment but irrespective of inoculum density and a competence development rate (X_B_, speed of propagation) correlated to the inoculum density (Figure 1D) ^39^. One hypothesis to explain the discrepancy between these studies was the presence of the polysaccharide capsule. We show that competence develops via propagation, irrespective of the presence of the capsule (Figure S1 and Supplementary Information) ^39,40^. We demonstrate that an accumulation of genetic and hardware factors prevented visualisation of the early stages of competence development in the previous study performed with the capsulated D39 strain (Figures S2-S4 and Supplementary Information) ^38^. To further investigate the propagative model of spontaneous competence, we observed competence development at the individual cell level, using a cell biology approach visualising fluorescent Cy3-DNA binding to transforming cells ^41^. This revealed propagation of competence throughout the population (Figures 1EF, S5), mirroring the theoretical propagation curves presented (Figure 1D). We further confirmed propagation of competence using sensitive single cell transformation assays (Figure S6 and Supplementary Information). Notably, the sensitivity of these assays revealed transformants prior to propagation, which may represent an initial subpopulation of self-inducing cells (Figure S6).

### Competence develops via Self-Induction and Propagation (SI&P) in response to antibiotic stress, shaping it as a populational health sensor

To further explore the existence of a self-inducing subpopulation at the origin of competence propagation, we repeated this experiment in non-permissive medium, tracking competence at the populational level via P*_ssbB_*::*luc* and at the cell level by sensitive transformation assays (Figure 2A). This revealed a time window where stochastic low-level self-induction was detected only at the cell level, but no populational propagation was observed (Figure 2B, top panels). In these conditions, competence is thus bimodal. As a control, we repeated this experiment with CSP added at 40 min, confirming that the ComABCDE QS system was able to respond in these conditions (Figure 2B, top panels). It was previously shown that antibiotics induce competence in these non-permissive conditions^33^. To explore competence propagation in these conditions, we conducted this experiment in the presence of sublethal concentrations of streptomycin. Competence propagation was observed after 150 minutes (Figure 2B, bottom panels), within the time window where self-induced cells were previously detected without propagation. To confirm that propagation developed within the entire population, this experiment was repeated with CSP added at 150 minutes, revealing competence induction with a steeper X_B_, showing that CSP addition artificially synchronised the population (Figure 2B, bottom panels). We deduced that in non-permissive medium, a stochastically self-induced subpopulation can be detected, but is insufficient to propagate competence throughout the population. Sublethal streptomycin addition, by increasing cellular stress levels, increases the self-inducing subpopulation, which reaches a threshold promoting competence propagation (Figure 2B, bottom panels). Similar observations were obtained using Mitomycin C (MMC) ^33^ (Figures S7 and S8 and Supplementary information). We unveil competence as a sensitive mechanism allowing individual cells to self-induce (SI) in response to stress, but only propagate (P) the signal in the face of sufficient stress. We call this mechanism Self-Induction and Propagation (SI&P). These experiments show that SI&P shapes pneumococcal competence as a population health sensor, able to evaluate environmental stress levels and react rapidly at the populational level.

**Figure 2:**
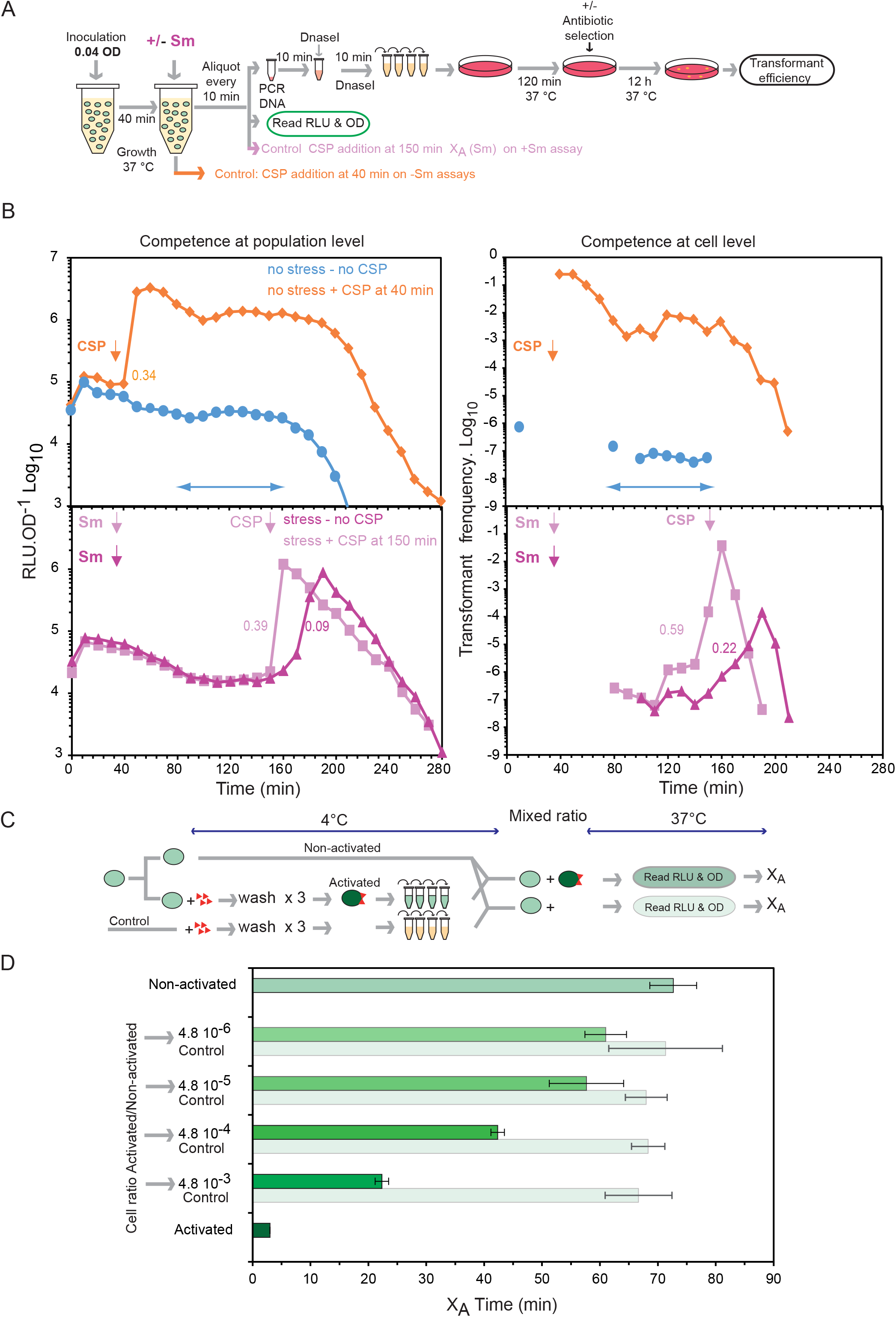
A self-induced subpopulation exists, and is increased upon antibiotic exposure, leading to propagation of competence throughout a population. (A) Schematic representation of experiment carried out to explore whether competence induction by exposure to sub-lethal concentrations of antibiotics follows an SI&P mode of transmission. (B) Competence levels tracked by RLU or transformation efficiency as shown in panel A. Left panels, RLU OD^-1^ tracking competence at the populational level; right panels, transformation efficiency tracking competence at the individual cell level. The blue arrowed lines highlight a period with detection of low transformant levels (right graph) revealing a self-induced cell fraction, without detection of populational competence propagation (left graph). Bottom panels show that antibiotic exposure increases this subpopulation, leading to competence propagation. Values represent rate calculations of competence development. Vertical arrows represent time of addition of CSP or antibiotic. Decrease in transformants and luminescence after ∼200 minutes are explained by entry into stationary phase (Figure S7A). Individual data shown representative of triplicate repeats showing similar results. (C) Schematic representation of experiment mixing activated and non-activated cells to artificially create fractions of competent cells and compare X_A_ times. Key as in Figure 1B. (D) A critical artificial cell fraction can initiate competence propagation. Full bars with a green decay correspond to the X_A_ values defined for varying ratios of activated/non-activated cells, framed by controls using 100% activated cells (dark green) and 100% non-activated cells (light green). Opaque light green bars correspond to the X_A_ values extracted from the negative control experiments (activated cells replaced with medium exposed to CSP). Standard deviations are reported. Individual data shown representative of triplicate repeats showing similar results.

### An artificially initiated competent cell fraction can propagate competence to non-competent cells

To explore the fraction of cells required to propagate competence, we attempted to identify the minimal artificial cell fraction able to propagate competence to non-competent cells. A wildtype non-competent cell culture was split in two and one sample was activated by exposure to CSP during 1 min and immediately washed at 4 °C to remove excess CSP not bound to the cells. The two samples were then mixed to generate varying ratios of activated and non-activated cells in permissive medium. The addition of artificially self-induced cells should shorten the X_A_ time, since the threshold of self-induced cells required for propagation will be reached more rapidly. The X_A_ time was compared to a control where CSP was added to cell-free culture, washed and used in the same ratio (Figure 2C). A population of only CSP-activated cells showed an X_A_ time of <5 min, consistent with immediate competence induction through artificial synchronisation (Figure 2D). In contrast, non-activated cells displayed an X_A_ of ∼70 min, resulting from spontaneous competence development. The mixed cultures showed a gradient of X_A_ values, revealing a clear correlation between cell ratio and X_A_ (Figure 2D). As the ratio decreased, the X_A_ increased until no clear effect can be attributed to the addition of activated cells. A difference was observed for the two least diluted ratios (4,8 10^-4^, 4,8 10^-3^), showing that the minimal self-inducing cell fraction required to promote populational competence is found between the 4,8 10^-4^ and 4,8 10^-5^ cell ratios. In these experiments, competence development results from a critical ratio of activated cells, mimicking the self-inducing cell fraction. This fraction can promote propagation of the competence signal through a population, further validating SI&P for the ComABCDE QS system.

### Competence improves tolerance to the genotoxic agents norfloxacin and methanemethylsulfonate

We have revealed that a growing pneumococcal culture is bimodal, with a minority of stressed, self-induced competent cells and a majority of non-stressed, non-competent cells. In our conditions, one self-inducing cell in 10^-5^ to 10^-6^ could be enough to propagate (Figure 2). We explored the benefits that inducing competence in non-stressed cells could provide. A previous study reported that competence improved survival of pneumococci upon transient exposure to three competence-inducing antibiotics ^23^. We further explored this competence-mediated benefit on cell survival by using two genotoxic drugs, norfloxacin (Norflo) and methanemethylsulfonate (MMS), which damage the replicating genome by blocking the gyrase or alkylating and depurinating nucleotides in DNA, respectively ^42–44^. Both induce competence of exponentially growing cells at sublethal concentrations ^33^ (Figure S9A).

To evaluate how survival of non-stressed, competent cells was impacted by these two stresses, we measured colony forming units (cfu) following exposure at above minimal inhibitory concentrations (MIC) (Figure S9BC and Table S1). We used strains which cannot naturally develop competence but can be induced by CSP, allowing rapid populational competence coordination. Cells activated (or not) during 10 minutes by CSP were exposed to above MIC concentrations of Norflo (100 µg mL^-1^) during 30 min or MMS (625 µg mL^-1^) during 15 min to compare survival (Figures 3A and S10A). Survival of non-competent cells reached 15 % and 1 %, respectively. By comparison, competent cells displayed improved survival, expressed as a tolerance ratio of 3,4 for Norflo and 4,3 for MMS (Figure 3B). These experiments elaborated on the previous finding ^23^ that competence increases tolerance of cells faced with drugs applied transiently above their MIC.

**Figure 3:**
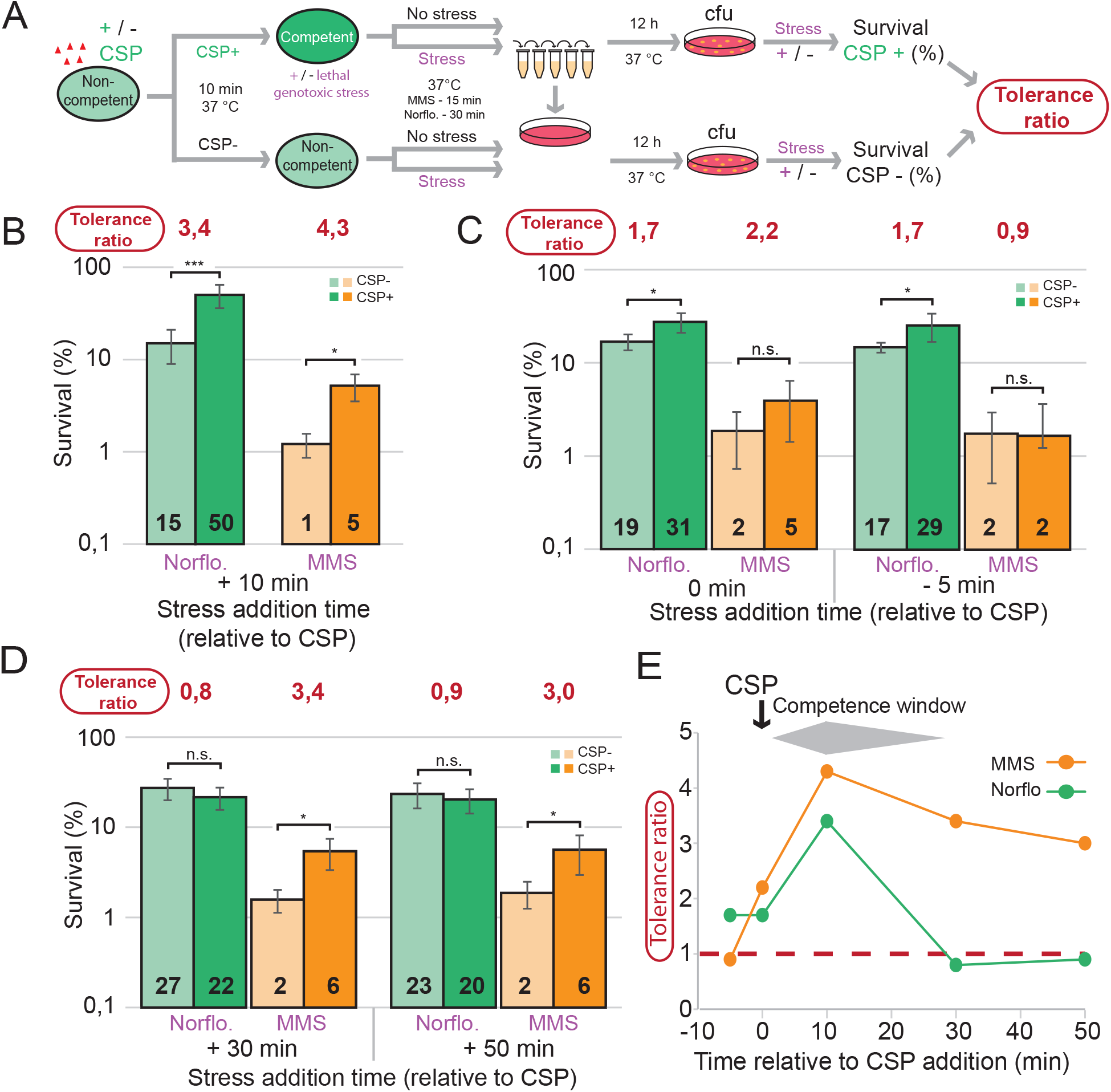
Increased survival of CSP-induced competent cells exposed to MMS or Norflo. (A) Simplified schematic of survival assay. Full experimental protocol in Figure S8. R1501 cells were induced to competence by CSP addition then exposed to stress. After 60 minutes, cells were serially diluted and plated. Tolerance ratios were calculated by comparing cfu of stressed and non-stressed populations. (B) Survival of competent (dark) and non-competent (light) cells exposed to Norflo or MMS at +10 min relative to CSP addition. Cells exposed to Norflo for 30 min and MMS for 15 min. Survival percentages calculated compared to non-stressed cells. Tolerance ratios calculated as in panel A. Experiments were carried out in triplicate with means and standard deviations presented. Asterisks represent significance between transformation efficiencies. *, p < 0,05; ***, p < 0,005. (C) Survival of competent and non-competent cells exposed to Norflo or MMS at –5 min or 0 min relative to CSP addition. Detailed assay description in Figure S8BC. Colour scheme, exposure times and representations as in panel B. n.s., non-significant, p > 0,05; *, p < 0,05; ***, p < 0,005. (D) Survival of competent and non-competent cells exposed to Norflo or MMS at +30 min and +50 min relative to CSP addition, representing post-competent populations. Colour scheme, exposure time and representations as in panel B. n.s., non-significant, p > 0,05; *, p < 0,05. (E) Tolerance ratio plotted against time of stress relative to CSP for Norflo and MMS. Data taken from panels B, C and D.

Next, we explored how the timing of stress exposure relative to CSP influences this improved tolerance. We added Norflo or MMS either 5 min before CSP or concurrently with CSP and measured the cfu following the same incubation time with these two drugs as above (Figure S10BC). Results showed that addition of CSP and stress concurrently reduced the tolerance ratio for Norflo and MMS (Figure 3C). Stress addition prior to CSP further reduced the tolerance ratio for MMS (0,9) but not Norflo. These findings show that the timing of stress exposure relative to competence induction modulates the beneficial effect of competence differently depending on the stress. To investigate whether the improved survival afforded to a pneumococcal population extended beyond the ∼30 minute competence window ^45^, we removed excess CSP 10 min after addition to prevent further cycles of competence (as can be observed with several peaks of transformation with CSP addition at 40 min in Figure 2B). Cells were then exposed to Norflo or MMS 30 or 50 minutes after CSP addition to assess survival of post-competent cells (Figure S10DE). Results showed that post-competent cells maintain increased tolerance to MMS, but not Norflo (Figure 3D). This further highlighted the heterogeneous effect of competence on descendent cell survival. This phenomenon can be visualised by plotting the tolerance ratio against the time of stress addition relative to CSP (Figure 3E).

Exposure to MMS for 15 min was sufficient to kill ∼99% of the population (Figure 3B), defining the minimal duration for killing 99 % of a population (MDK_99_), a value used as a metric of bacterial tolerance ^46,47^. In contrast, the MDK_99_ was not established for Norflo as survival remained around 15% (Figure 3B). To establish the MDK_99_ for Norflo, a time course was carried out with survival of non-competent and competent cells compared at varying time-points after Norflo addition. MDK_99_ values of ∼74 and ∼103 min exposure were respectively determined (Figure S9D). In conclusion, competent cells display improved tolerance to the genotoxic stresses MMS and Norflo. In addition, tolerance ratios are modulated dependent on when stress addition occurs compared to competent induction, highlighting a heterogeneous behaviour of competent cells in overcoming these stresses.

### Competence modulates survival of cells exposed to a wide variety of lethal stresses

To further investigate tolerance of competent cells faced with lethal stresses, we enlarged the analysis to different antibiotics, targeting basic cellular functions: cell wall synthesis (ampicillin, vancomycin), genome integrity (mitomycin C, novobiocin, trimethoprim), transcription (rifampicin) and translation (erythromycin, tetracycline, streptomycin, kanamycin). Some of these drugs induce competence when applied at sub-MIC concentrations (Table S1). For ten of twelve stresses applied above their MIC (Figure S11) for 60 min, competent cells displayed increased survival compared to non-competent cells, with tolerance ratios between 2,5 and 7,8 (Figure 4A). By stark contrast, competent cells displayed increased sensitivity to aminoglycosides (streptomycin and kanamycin), with tolerance ratios of 0,0024 and 0,04 respectively (Figure 4A). Aminoglycosides were previously found to induce competence ^33^. Since these effects were observed at a single time point, time course experiments were carried out for stresses for which competence increases survival (vancomycin) or sensitivity (kanamycin), which showed that these effects were gradual between 20 and 60 minutes post-stress exposure (Figure S12AB), as for norfloxacin (Figure S9D). In conclusion, competence modulates the survival of cells transiently exposed to a wide variety of lethal stresses, with a general benefit for competent cells, revealing phenotypic heterogeneity within the competent population.

**Figure 4:**
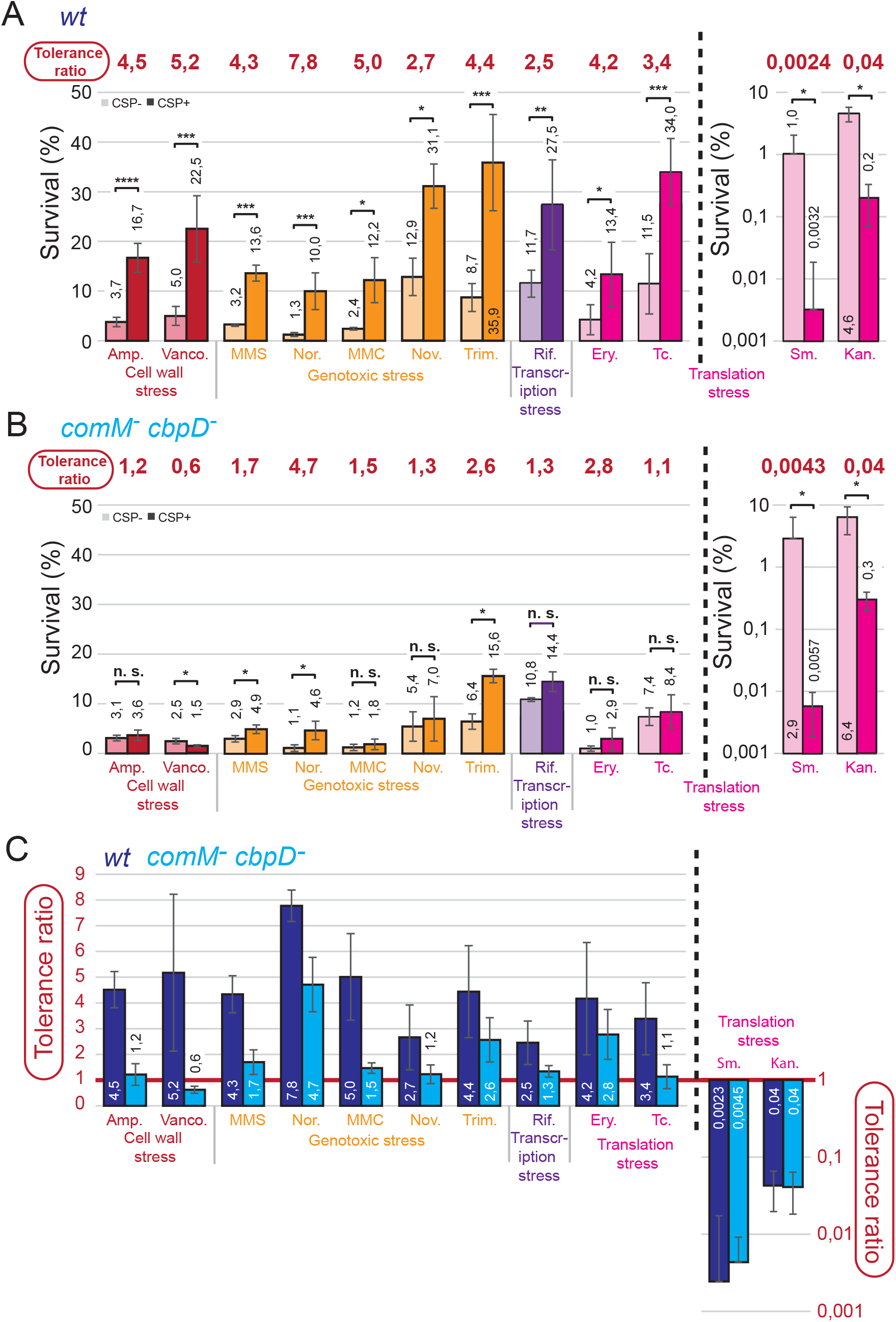
Competence modulates survival of pneumococci exposed to a wide range of stresses, with ComM key to increased survival. (A) Survival of competent (dark) and non-competent (light) R3369 cells exposed to various stresses for 60 min starting at +10 min relative to CSP addition. Full experimental protocol in Figure S8A. Colours represent different types of stress. Black dotted line separates stresses where competence increased survival from those where competence increased susceptibility, plotted on a secondary y axis. Tolerance ratios calculated as in Figure 3A. Experiments were carried out at least in triplicate with means and standard deviations presented. Asterisks represent significance between survival of competent and non-competent cells. *, p < 0,05; **, p < 0,01; ***, p < 0,005; ****, p < 0,001. (B) Survival of competent (dark) and non-competent (light) *comM^-^ cbpD^-^* cells (R4592) exposed to various stresses for 60 min starting at +10 min relative to CSP addition. Experimental procedures and representations as in panel A. n.s., non-significant, p > 0,05; *, p < 0,05. (C) Comparison of tolerance ratios of wildtype and *comM^-^ cbpD^-^* cells, with values of 1 representing no difference between competent and non-competent cells, values >1 representing increased survival in competent cells and values <1 representing increased survival in non-competent cells.

### ComM is central to the improved tolerance to stress of competent cells

Increased tolerance has been associated with reduced growth and altered metabolism in other bacteria ^48–52^ could play a role in competence-mediated tolerance increase. To test this hypothesis we inactivated *comM* (and the hydrolase *cbpD*, to which ComM provides immunity ^14^), and repeated the screen with the isogenic wildtype strain. The absence of *comM* reduced the tolerance ratio for stresses where competence had increased survival, revealing ComM as key to increased survival during competence (Figure 4BC). In some cases, the tolerance ratio was reduced to ∼1, suggesting that ComM alone was responsible for the increased survival during competence, but in others a tolerance ratio of >1 remained, suggesting other competence factors may also contribute. These results further highlight the heterogeneity present in a competent pneumococcal population. Similar tolerance ratios were observed between control strains lacking only *comM* or *cbpD* (Figure S12CDE), showing that the observed effect was mediated by ComM alone. In contrast, absence of *comM* (and *cbpD*) did not alter the tolerance ratio for the two aminoglycoside stresses where competence increased sensitivity (Figure 4BC). In conclusion, competence-mediated tolerance increase is mainly dependent on the early competence protein ComM.

### Competent-tolerant cells able to transform favour acquisition of gene cassettes over single nucleotide polymorphisms

We defined competent cells that tolerated stress as competent-tolerant and explored whether these cells retained the ability to transform under lethal stress. We used tDNA fragments with a selective antibiotic resistance provided via a heterologous gene cassette (HGC) or a single nucleotide polymorphism (SNP). We performed the survival assay with stress added 10 min after CSP, and exogenous transforming DNA (tDNA) was added at the same time as the stress, followed by 60 min exposure in liquid medium. Next, cells were diluted, plated and incubated for 80 min before transformant selection. This procedure allowed determination of transformation efficiency, along with survival and tolerance ratios (Figure 5A and S13). In HGC assays performed under five different stresses targeting different cell processes, the transformation frequency of tDNA conferring kanamycin resistance was reduced 3-8-fold, in comparison to a non-stressed population (Figure 5B). In stark contrast, in the SNP assay using tDNA containing an *rpsL41* point mutation (A157C transversion) conferring streptomycin resistance ^53,54^, a significantly larger decrease in transformation efficiency was observed, ranging from 43-156-fold (Figure 5C). Thus, although the transformation efficiency of SNPs is 50-fold higher than that of HGC under non-stressed conditions, the former was much more affected by stress. This was also the case with another SNP in the *rpoB* gene (C1517T transition), conferring rifampicin resistance, *rif23* ^53^ (Figure 5D). Thus, transformation efficiency was altered in competent-tolerant cells, as fewer cells were able to transform, further highlighting the adaptive heterogeneity of a competent population exposed to stress. The Hex mismatch repair (MMR) system interferes with transformation at the recombination step by recognising mismatches generated during heteroduplex formation. MMR recognises mismatches generated by transition (*rif23*) better than those generated by transversion (*rpsL41*) ^55^, explaining the ∼4-fold deficit in transformation between these markers in non-stressed cells (Figure 5CD). We repeated the assay in an isogenic MMR^-^ strain. As expected, the same transformation efficiency was observed for *rif23* and *rpsL41* in the absence of stress. Interestingly, we observed increased transformation efficiency of these two SNPs in competent-tolerant MMR^-^ cells compared to MMR^+^ cells (expressed as MMR ratio; Figure 5E). These results strongly suggest that MMR is hyper-active in competent-tolerant cells. In conclusion, competent-tolerant cells transform at low levels and favour HGC over SNP transfer due to increased MMR.

**Figure 5:**
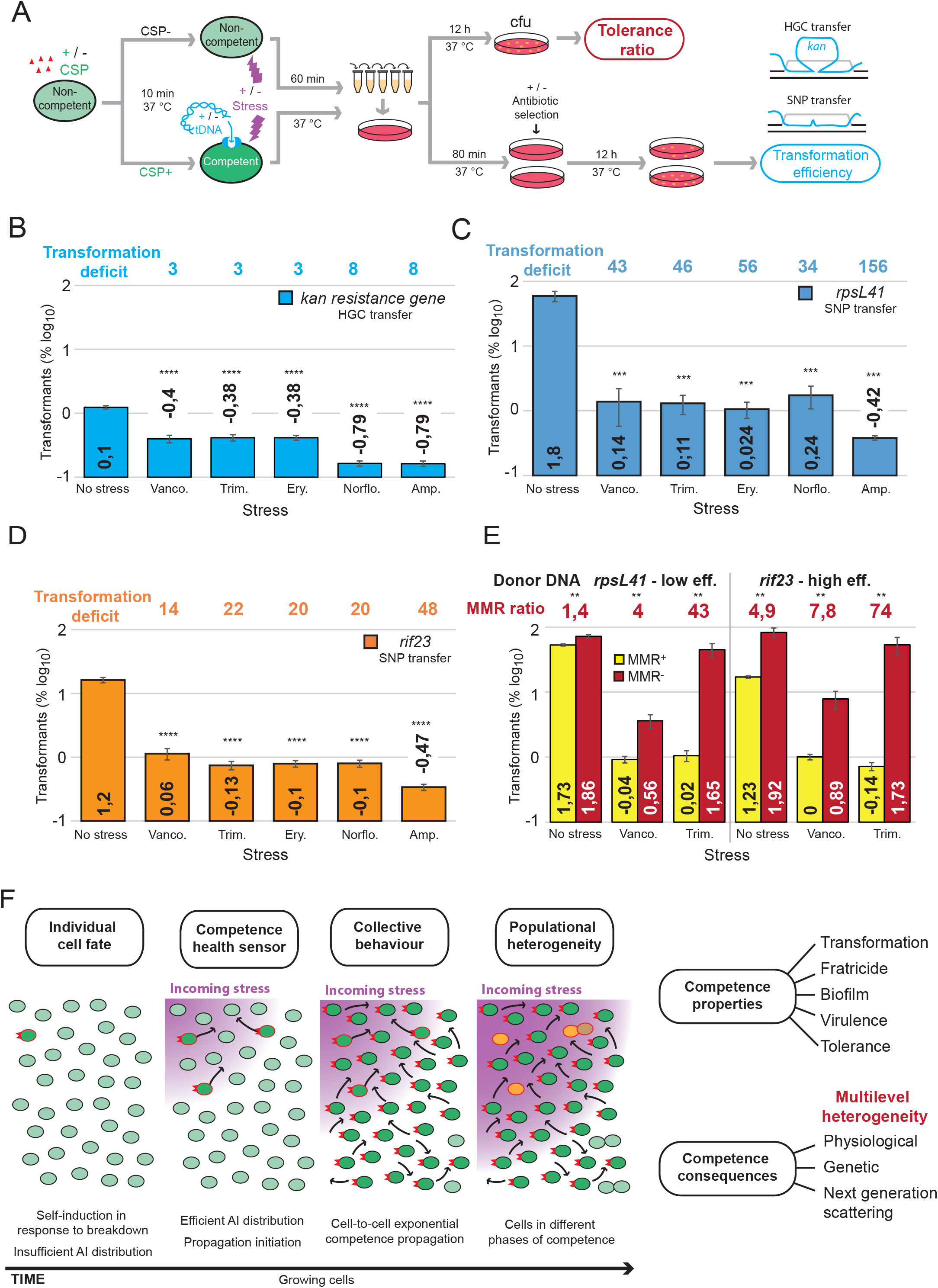
Tolerant cells favour integration of heterologous cassettes over point mutations by transformation. (A) Simplified schematic of transformation survival assay. Full experimental protocol in Figure S13. Cells were induced to competence by CSP addition then exposed to stress and/or tDNA 10 min later. After 60 minutes, cells were serially diluted and plated, with a second layer added for antibiotic selection 80 minutes later. Tolerance ratios were calculated by comparing cfus of stressed and non-stressed populations. Transformation efficiency was calculated by comparing cfu on selective and non-selective plates. Transformation diagrams represent formation of a D-loop structure for integration of a HGC or SNP flanked by homologous sequence. Black lines represent recipient chromosome, with displaced DNA in gray and tDNA in blue. (B) Transformation efficiency of heterologous sequences in tolerant pneumococcal cells. Protocol carried out as in panel A, with cells transformed with tDNA possessing a heterologous cassette conferring kanamycin resistance (*comFA::kan)*. Diagram represents formation of a D-loop structure for integration of the heterologous *kan* sequence in the *comFA* locus. Black lines represent recipient chromosome, with displaced DNA in gray and tDNA in blue. Experiments were carried out in triplicate with means and standard deviations presented. Asterisks represent significance between transformation efficiencies. ****, p < 0,0001. (C) Transformation efficiency of *rpsL41* SNP in tolerant pneumococcal cells. Transformation survival assay carried out and tolerance ratios calculated as in Figure S13. Cells transformed with tDNA possessing SNP conferring streptomycin resistance (*rpsL41*). Experiments were carried out in triplicate with means and standard deviations presented. Asterisks represent significance between transformation efficiencies. ***, p < 0,005. (D) Transformation efficiency of *rif23* SNP in tolerant pneumococcal cells. Transformation survival assay carried out and tolerance ratios calculated as in Figure S13. Cells transformed with tDNA possessing SNP conferring rifampicin resistance (*rif23*). Experiments were carried out in triplicate with means and standard deviations presented. Asterisks represent significance between transformation efficiencies. ****, p < 0,0001. (E) Comparison of transformation efficiencies of point mutations (*rpsL41* and *rif23*) in MMR+ and MMR^-^ cells. Transformation survival assay carried out as in panel *A*. Experiments were carried out in triplicate with means and standard deviations presented. Asterisks represent significant difference between MMR^+^ and MMR^-^ populations (MMR ratio). **, p < 0,001. (F) Schematic representation of competence development. At low environmental stress levels, a small fraction of cells self-induces competence but does not distribute sufficient CSP to propagate competence. As stress increases, the self-inducing fraction reaches a threshold of CSP distribution able to convert neighbouring cells to competence. This reveals competence as a populational health sensor. Competence can propagate exponentially rapidly through a population, ensuring that cells not sufficiently stressed to self-induce become competent prior to stress exposure. This wave of competence generates populational heterogeneity, with a mixture of competent and post-competent cells. Competence provides a population with several properties, as well as promoting heterogeneity at multiple levels. This includes physiologically by altering tolerance to stress, genetically via transformation, and within the next generation via lop-sided transmission of DprA, affecting subsequent competence waves. Cell identities as in Figure 1B, with different shades of orange representing lop-sided DprA transmission to daughter cells. Purple wave represents incoming environmental stress. Black lines represent propagation of competence throughout the population.

## Discussion

This work validates propagation as the general mode of pneumococcal competence transmission and reveals individual self-induced cells as the source of populational competence development. We call this mode SI&P (Figure 2). Here, we present arguments showing that SI&P shapes competence as a populational health sensing mechanism equipped to discriminate incoming stress from false alarm. Competence relies on stochastic self-induction of a minority of cells in response to stress, while the majority are non-stressed but induced via propagation (Figure 5F). In transformation assays, we detected one self-induced cell per 10^-7^, but competence propagation required a greater ratio (Figure 2). The planktonic growth condition governs the probability of cell-to-cell contact and we deduce that the ratio of CSP-decorated self-induced cells to non-stressed cells must pass a threshold to propagate and convert the non-stressed cells (Figures 2 and 5F). The role of unphosphorylated ComE as a transcriptional repressor should contribute as a mechanism guarding against false alarm, with the stoichiometry between ComE and the transcriptional activator ComE∼P governing competence development (Martin et al., 2013) and generating a populational heath sensor. Once CSP distribution by cell-to-cell contact overcomes this protective mechanism, competence propagates. Such a mechanism benefits the population since it can avoid the cost of populational competence development via false alarm but react efficiently to a stressful environment, rapidly converting the population to competence (Figure 2). Propagation is exponential since the AI is retained on the cell surface, unlike AI diffusion, which leads to a linear response. Noise and stochasticity are key to the appearance of the SI fraction, but additional environmental stresses increase this fraction, allowing the cell population to discriminate between noise and incoming stress (Figure 2). SI has been observed in two QS systems in *Bacillus subtilis* as well as in *Pseudomonas aeruginosa* (Bareia et al., 2018; Smith and Schuster, 2022). However, no populational propagation was observed in these systems. Altogether, these results reveal pneumococcal competence as a health sensor of the population, idling in the absence of danger but able to react extremely rapidly when danger is detected.

Pneumococcal competence not only acts as a populational health sensor, but also generates multilevel heterogeneity within a population as revealed by altered ability to tolerate transient exposure to lethal stresses. Competence increases tolerance to most tested stresses, except aminoglycosides, for which it dramatically increases sensitivity (Figure 4). Improved tolerance was observed for various lethal stresses targeting vital cellular functions (genome replication, translation, peptidoglycan synthesis…), highlighting the wide-ranging protective effect provided by SI&P driven competence. Improved tolerance against lethal drugs was also reported for competent *B. subtilis* cells, linked to ComGA-mediated growth arrest ^51^. We show that improved pneumococcal tolerance also mainly depends on a transient growth delay mediated by the early competence protein ComM ^15^. Nonetheless, competent cells lacking ComM still exhibit improved tolerance to some stresses (Figure 4C), showing that other competence-induced effectors provide specific protection. Remarkably, there is no relation between increased tolerance to a particular stress and its ability to induce competence, suggesting that competence induction diversifies a population, providing the potential to tolerate a wide variety of stresses. Competence is known to provide transformation ability to all cells, potentially scattering a clonal population into a mosaic of genotypes. However, under lethal stress, competent-tolerant cells are biased towards transformation-mediated acquisition of HGCs over SNPs, due to increased MMR activity. Another level of heterogeneity comes via DprA, which accumulates at one cell pole to promote competence shut-off ^56^. This results in lop-sided inheritance of DprA in daughter cells, which partly explains the poor coordination of secondary competence waves ^28,29^. Overall, this shows that the competence health sensor fosters functional diversification via multilevel heterogeneity of a clonal population (genetic heredity, physiological state, physiological heredity) (Figure 1C and 5F), with some cells better suited to surviving various stresses.

Pneumococcal competence development not only promotes multilevel heterogeneity, but self-induction is also driven by stochastic diversity between members of the population. This diversity generates a bimodality, with a minority of cells self-inducing and the majority remaining non-competent unless propagation conditions are met. This bimodality is a common feature of QS systems in other bacterial species, such as *B. subtilis*, where competence is induced in a fraction of cells based on the noisy expression of its master transcriptional regulator ComK ^57,58^. *Streptococcus salivarius* also displays bimodality in competence development. The ComR transcriptional regulator ^59^ is negatively controlled by a two-component regulatory system, CovRS, with bimodality coming from heterogeneity in sensing environmental changes, without observed propagation ^60^. A key difference compared to these two systems is that the bimodality of pneumococcal competence can lead to populational propagation. Bimodal competence development has been suggested as a bet hedging strategy ^60,61^ where only a fraction of the population pays the energetic cost of competence but also reaps the potential benefits. This cannot be applied to pneumococcal competence due to its propagative and populational nature. Nonetheless, populational competence development, which initially depends on bimodality, scatters a population towards multilevel heterogeneity, fitting the definition of bet hedging. In contrast to both *B. subtilis* and *S. salivarius*, the pneumococcus is an important opportunistic human pathogen ^62^, exposed to environmental stresses such as host immunity and antibiotics during both commensal and pathogenic life. Our work shows that competence is tightly integrated into the pneumococcal lifestyle and key for the survival of this major human pathogen. Competence-mediated diversification of a clonal pneumococcal population may provide the best opportunity for survival in such environments. This idea is strengthened by the links between pneumococcal competence and virulence ^18,20,22,63^, and the suggestion that a single cell bottleneck is at the origin of pneumococcal bacteraemia ^64^.

This study reveals that pneumococcal competence is a health sensor responding to phenotypic heterogeneity and proceeding via an SI&P mode in the face of environmental stresses (Figure 5F). Competence development drives multilevel heterogeneity, with one result being altered tolerance in the face of transient lethal stress exposure. Competence can thus be framed as a global pneumococcal stress response, with transformation representing one of the multiple facets of this response. In conclusion, SI&P shapes pneumococcal competence as a health sensor tuned to scatter the clonal population towards multilevel diversification, employing a bet hedging strategy to maximise the survival potential of a pneumococcal population in response to environmental stresses such as antibiotics.

## Materials and Methods

### Bacterial strains, construction and transformation

*S. pneumoniae* strains and primers used in this study are described in Table S3. CSP-induced transformation was performed as described previously ^65^ using pre-competent cells treated at 37 °C for 10 min with synthetic CSP1 (100 ng mL^-1^). After addition of tDNA, cells were incubated for 20 min at 30 °C. Transformants were selected by plating in 10 mL CAT-agar supplemented with 4 % horse blood, incubating for 2 h at 37 °C for phenotypic expression, and overlaid with 10 mL CAT-agar containing the appropriate antibiotic as followed: chloramphenicol (4.5 µg mL^-1^), kanamycin (250 µg mL^-1^), streptomycin (200 µg mL^-1^), tetracycline (1 µg mL^-1^). To generate strain R2287, R1501 was transformed with a mariner mutagenesis fragment of the *comFA* gene, generated using primer pair comFA3-comFA4, with the pR410 plasmid, as previously described ^66^. A *comFA::kan* transformant with a kanamycin resistance cassette inserted in the *comFA* gene in a co-transcribed orientation was isolated by selection with kanamycin. To generate the R4428 strain, a kanamycin allele knock-out with a 7 bp deletion in the ORF was generated using the primer pairs MP259-MP2260 and MP261-MP262 to amplify two DNA fragments from strain R2737, followed by splicing by overlap extension (SOE) PCR using primer pair MP259-MP262. This DNA fragment was transformed into R2737 ^67^, and a kanamycin-sensitive transformant was isolated. The presence of the *kan^S^*allele was validated by sequencing. To generate strain R4590, R3369 (*comC2D1*) was transformed with genomic DNA (gDNA) from strain R3967 ^15^, and *comM::cat* transformants were isolated by selection with chloramphenicol. To generate strain R4591, R3369 (*comC2D1*) was transformed gDNA from strain R1620 ^68^, and *cbpD::spc* transformants were isolated by selection with spectinomycin. To generate strain R4592, R4590 (*comC2D1, comM::cat*) was transformed with gDNA from strain R1620 ^68^, and *cbpD::spc* transformants were isolated by selection with spectinomycin. To generate TD277, D39_Tlse_ was transformed with gDNA from strain R3316 ^36^, with P*_ssbB_*::l*uc* transformants isolated by selection with chloramphenicol. To generate strain TD288, D39V ^69^ was transformed with gDNA from strain R3316 ^36^, with P*_ssbB_*::l*uc* transformants isolated by selection with chloramphenicol.

### Spontaneous competence detection using luciferase gene reporter

Competence development was monitored using P*_ssbB_*::*luc* as described previously ^40^. In these experiments, C+Y medium at different initial pH ranges was used to control the ability of cells to spontaneously develop competence, as described previously ^39^, with acidic pH 6,8-7,0 being ‘non-permissive’ to competence and alkaline pH 7,6-7,9 ‘permissive’. This nomenclature will be used throughout this manuscript. Briefly, cells were grown for several generations to mid-log phase in non-permissive medium, washed and stored at -70 °C in fresh medium supplemented with 20 % glycerol. This cell stock was used to generate varying inoculum sizes in permissive medium in a clear-bottomed 96-well white NBS microplate (Corning). Cells were grown at 37 °C in a Varioskan Flash (Thermo 399 Electron Corporation) luminometer and relative luminescence units (RLU) and OD values were recorded throughout incubation.

### Spontaneous competence detection using transformation assays

14 mL of permissive medium was inoculated with two different cell densities at 37°C to provide high and moderate X_B_ competence development rates ^39^. During growth, every 10 minutes, 100 µL of cells were tested for transformation after 10- or 20-minutes exposure to tDNA and 300 µL were used to measure photon emission and OD. A control was generated by the addition of synthetic CSP after 70 minutes of growth to obtain an artificial fully coordinated competent population ^45,70^. Each transformation assay was carried out in 100 µL aliquots of cell culture. tDNAs used for measuring transformation efficiency were the purified 3,434 bp PCR fragment generated with the primer pair MB117-MB120 containing a SNP conferring streptomycin resistance ^71^ or the purified 3,238 bp PCR fragment containing the wildtype kanamycin resistance allele generated by the primer pair MP259-MP262. Both tDNA PCR fragments were used at 200 ng mL^-1^. The streptomycin resistance allele differs by a single nucleotide from the sensitive wildtype allele. The kanamycin sensitive allele differs from the kanamycin resistant allele by a 7-nucleotide deletion in the open reading frame. Both mutations are central within the PCR fragments used. Without transformation, no spontaneous kanamycin resistant clones were detected in R4428. Cells are incubated with tDNA at 30 °C during 10 or 20 min followed by addition of 20 µg mL^-1^ DNase I (Sigma Aldrich) with MgCl_2_ at 10 mM for a further 10 minutes at 37 °C before plating. Selection of the transformants was done using streptomycin at 200 µg mL^-1^ or kanamycin at 250 µg mL^-1^. Controls with no tDNA or with concomitant addition of tDNA and DNase I were repeated several times and gave no detectable transformants.

### Spontaneous competence detection by visualisation of DNA binding using microscopy

Analysis of DNA binding was performed in a virulent *endA* mutant (TD290), to favour accumulation of transforming DNA at the surface of competent cells. In wildtype, *endA^+^* cells, surface-bound DNA is immediately internalised into the cytosol or degraded, which makes surface-bound DNA accumulation hard to visualise ^41^. Precultures were inoculated at two cell densities (8 10^-3^ and 8 10^-4^ OD_550_) in permissive medium and allowed to grow, with samples taken at various time points through growth to determine DNA binding. X_A_ detection was carried out by using P*_ssbB_*::*luc*. The positive control with CSP addition on the low cell density inoculum was carried out at the X_A_ time. For each time point along growth an aliquot was incubated for 5 min with 10 ng of a 285 bp DNA fragment labelled with a Cy3 fluorophore at its 5’ extremity, amplified using primer pair OCN75-76 ^41^. Cells were pelleted (3000 x g, 3 min), washed twice in 500 µL C+Y, and resuspended in 20 to 50 µL C+Y medium before microscopy. Two µL of this suspension was spotted on a microscope slide containing a slab of 1.2% C+Y agarose as described previously ^72^. Phase contrast and fluorescence microscopy were performed with an automated inverted epifluorescence microscope Nikon Ti-E/B, a phase contrast objective (CFI Plan Apo Lambda DM 100X, NA1.45), a Semrock filter set for Cy3 (Ex: 531BP40; DM: 562; Em: 593BP40), a LED light source (Spectra X Light Engine, Lumencor), and a sCMOS camera (Neo sCMOS, Andor). Images were captured and processed using the Nis-Elements AR software (Nikon). Cy3 fluorescence images were false coloured red and overlaid on phase contrast images. Overlaid images were further analysed to quantify the number of cells bound with Cy3-labelled DNA. Single cells were first detected using the threshold command from Nis Elements and cells bound or not to DNA were manually classified using the taxonomy tool. Data representative of three independent repeats.

### Cell mixing experiment

A stock of the R4428 strain grown in non-permissive medium conserved in glycerol at -70 °C was used to inoculate non-permissive medium at 37°C at OD_550_ 0,004. Cells were grown to OD_550_ 0,06 and growth was stopped on ice. Cells were washed by centrifugation and resuspended in permissive medium at 4 °C. A cell aliquot and a sterile medium aliquot without cells (negative control) were exposed to 25 ng mL^-1^ of CSP during one minute at 37 °C then placed on ice. Each assay was washed 3 times by centrifugation with the same volume of sterile permissive medium at 4 °C. Different dilutions of both cell aliquot and sterile medium aliquots were mixed with a constant number of cells not exposed to CSP and loaded directly on a clear-bottomed 96-well white NBS micro plate (Corning). The microplate was incubated at 37°C immediately and the Relative luminescence units (RLU) and OD recorded every minute in a Varioskan Flash (Thermo 399 Electron Corporation) luminometer. Cfu were counted to obtain the ratio in the assays between cells exposed (or not) to CSP.

### Survival assays

Strains used for these experiments were unable to spontaneously induce competence since they either possessed a *comC0* mutation ^11^ or the combination of the non-compatible alleles *comC_2_* and *comD_1_* ^18^. The experiments are summarised in Figure S10. Strains were grown in permissive medium to OD_550_ 0,2, washed and concentrated to OD_550_ 0,4 in C+Y + 20 % glycerol and stored at -70°C. Cells used to inoculate were either defrosted or pre-inoculated and grown exponentially to OD_550_ 0,1, followed by dilution in permissive medium at OD 0,004 and incubation at 37 °C. Cells were grown during 20 min at 37 °C and divided into two samples, one with 100 ng mL^-1^ CSP, the other sample without. Each sample was incubated for a further 10 min and again divided into two samples, one exposed to stress and one not. After 60 min incubation at 37 °C, cells were serially diluted, plated at appropriate dilutions on CAT-agar supplemented with 4% horse blood (two plates per dilution) and incubated at 37 °C overnight. 60 min was chosen as a stress exposure time since this was close to the MDK_99_ of norfloxacin (Figure S9). Other timings used are as described in the text. For time-course experiments, culture volumes were scaled up, and cfu were measured for each time point. Stress concentrations used were above the MIC and can be found in Table S1. Comparison of total cfu between cultures exposed or not to stress allowed calculation of survival as a percentage, with comparison of competent and non-competent percentages showing the effect of competence on survival. Each experiment was repeated at least 3 times. Mean and standard deviations were calculated for each experiment, and a Student’s t-test was used to determine the significance of the difference in survival between competent and non-competent populations.

### Transformation survival assays

Transformation survival assays were carried out based on survival assays used in this study, but with modifications as shown in Figure S13. Briefly, precultured cells were rediluted in fresh medium at OD_550_ 0,004 and incubated for 20 min at 37 °C before being split into two samples, one with 100 ng mL^-1^ CSP, the other sample without CSP. Samples were then incubated for 10 min at 37°C. Non-competent cells were then split in two, one exposed to stress and one not. Competent cells were split into four, with and without stress and tDNA. After 60 min incubation at 37 °C, cells were serially diluted and plated at appropriate dilutions to detect total cfu and transformant cfu on CAT-agar supplemented with 4% horse blood (two plates per dilution). Plates were incubated at 37 °C for 80 min to allow phenotypic expression of transformed phenotypes, before a second layer of medium containing appropriate antibiotic was added to transformant cfu plates for selection. Plates were then incubated overnight at 37°C. Transformation efficiencies were calculated by comparing total cfu with transformant cfu. Stress concentrations used can be found in Table S1. Three DNA fragments were used for transformation, an *rpsL41* point mutant conferring streptomycin resistance, a *rif23* point mutant conferring rifampicin resistance, and a *comFA::kan* cassette conferring kanamycin resistance. A 3,434 bp DNA fragment containing *rpsL41* was amplified from strain R304 using primer pair MB117-MB120. A 4,195 bp DNA fragment containing the *rif23* point mutation was amplified from strain R304 using primer pair MB137-MB138. A 4,931 bp DNA fragment containing *comFA::kan^3C^* was amplified from strain R2287 using primer pair CJ339-CJ356. Antibiotics used for selection were streptomycin (200 µg mL^-1^), rifampicin (2 µg mL^-1^) and kanamycin (250 µg mL^-1^). Each experiment was repeated in triplicate, with means and standard deviations calculated, and a Student’s t-test used to determine the significance of the difference in transformation efficiencies.

## Acknowledgements

We thank Jan-Willem Veening for providing us with strains and medium, allowing us to understand the differences between the two studies.

## Supplementary information

### Introduction

In this section, we report evidence demonstrating that spontaneous pneumococcal competence development relies on a propagative mechanism, whatever the genotype of the cells. In all conditions tested, the two key parameters X_A_ and X_B_, which report the development time and the rate of pneumococcal population development during growth, respectively, fit with a propagative mechanism that we previously described ^39^ (Figure 1CD). In this mechanism, the X_A_ value is independent of the cell density, while the X_B_ value is linked to cell density. We previously defined this propagative mechanism by recording the luciferase activity of a transcriptional fusion between the luciferase gene with the promoter of the late competence gene *ssbB* (P*_ssbB_*::*luc*) in an unencapsulated strain. Using the same approach with the capsulated D39 strain, Moreno and colleagues reported a different mechanism of competence development akin to classical QS, where the X_A_ value varied in function of the cell density of the inoculum and the X_B_ value remained constant ^38^. Here, we explain the differences observed between these two studies and confirm propagation as a general mechanism of pneumococcal competence development.

### Results and discussion

#### Spontaneous competence development in the population proceeds by propagation, independent of polysaccharide capsule presence or growth medium

Considering the reported differences between these two studies, we reproduced these experiments using the same methods and confirmed that pneumococcal competence development proceeds via propagation, irrespective of the presence of the polysaccharide capsule (Figure S1). Another possible difference between the studies could come from environmental growth conditions including medium used. To remove all doubt, we tested C+Y medium kindly provided by the Veening laboratory. The experiment was repeated with both media made in Toulouse (Tlse’s C+Y medium) and Lausanne (Lsne’s C+Y medium). Notably, both laboratory media led to comparable propagative behaviour for competence development (Figure S2A), showing that medium differences did not explain the different results observed.

#### Confirmation of propagation as the general mechanism of pneumococcal competence development

In these experiments, we observed differences in readings during the X_A_ time (Figures S1 and S2), which could affect detection of initial competence development, and could be attributable to differences in genotype, reporter construction and/or sensitivity of the luminometer. We investigated these hypotheses individually. First, a 5-fold decrease in basal competence level was observed in the D39_(Tlse)_ lineage compared to its R800 lineage derivative, irrespective of the presence of the capsule (Figure S1A). We attribute these difference to the genetic background, since minor differences can modify the fine-tuning of the idling ComABCDE QS system ^39^. We also observed differences in basal expression levels between D39_(Tlse)_ and D39_(Lsne)_ (Figure S2A), which possess P*_ssbB_*::*luc* reporter fusions located at native or ectopic loci respectively. Exploration of this demonstrated that the ectopic P*_ssbB_*::*luc* reporter was less sensitive than the native one, leading to an approximate 50-fold loss of sensitivity compared to R800 (Figure S2A). This renders the reported values prior to competence induction too close to the limits of detection for cell densities where they are readily detectable for the P_ssbB_::*luc* at *ssbB* reporter (Figure S2B, red circles). In addition, by comparing the sensitivity of the luminometers used in both studies to a reference luminometer ^40^, we revealed a 16-fold deficit in sensitivity for the Tecan luminometer used by Moreno and colleagues (Figure S3). To conclude, precise and exhaustive dissection of the tools used to report competence has explained the differences observed between the two studies ^38,39^. Altogether, these results confirm that propagation is the general mechanism of competence development in pneumococci, as previously proposed ^39^.

#### Dilution of competent cells does not abolish competence

A previous study suggested that dilution of competent cells rendered these cells non-competent ^38^. We hypothesised that if competent cells were diluted, the competence status should not be modified. To explore this, we repeated the experiments conducted in the study ^38^, and included a transformation assay as a second independent method to report competence. Whatever the competence state reported by P*_ssb_*::*luc* RLU, if a cell fraction is competent, this should be detected in a transformation assay ^40^. We focussed our analysis on the first minutes of the culture. As expected, the first recorded RLU values diminished by tenfold between each assay, which correlates to the reduction of the inoculum sizes, except for the two lowest inocula (Figure S4A, red circles) explained by the sensitivity limit of the luminometer (Figure S3). The specific RLU OD^-1^ activity calculated on the first recorded data confirms that dilution until 10^-4^ OD gives equivalent values for the cell population in competence recording (Figure S4B, blue numbers, left-graph). In parallel, the transformation assay gives a transformant ratio approximately fitting the theoretical transformation efficiencies estimated by the serial dilutions (Figure S4B, right panel, red dotted line). Thus, dilution of a competent population does not render cells non-competent. In addition, we repeated the mirror experiments but using cells precultured in acidic medium (non-permissive) (Figure S4C). We can approximately plot the competence threshold rule (200 RLU) from the previous study ^38^ transposed to the RLU produced by our luminometer (Figure S4C medium panel, red dotted line). By doing so we are able to show that the conditions previously used ^38^ prevented visualisation of an unmodified X_A_ whatever the inoculation density but a X_B_ linked to the cell density (Figure S4C).

#### Competence propagation visualised by fluorescence microscopy

To observe competence propagation using microscopy (Figure 1E), an *endA* mutant, in which fluorescent Cy3-tDNA accumulates at the septum of competent cells, was used ^41^. High- and low-density cultures of *endA* mutant cells possessing P*_ssbB_*::*luc* were grown in permissive medium for competence to calculate the X_A_ time in these conditions (Figure S5A). Then, samples were taken every 10 min after this time, and Cy3-tDNA added, before visualisation on an agar pad (Figures 1F and S5B). A control sample of low cell density inoculum with CSP added at the X_A_ time was included. Results show an exponential increase in the frequency of cells binding Cy3-tDNA (Figure 1F), from which the competence development rate can be calculated. The rate observed was significantly slower in the low-density inoculum. These results reveal propagation of competence with a rate corresponding to an exponential function that depends on cell density ^39^.

#### Sensitive transformation assays validate propagation which may originate from self-inducing competent cells, establishing bimodal behaviour

We used transformation assays to further demonstrate propagation. We compared transformation levels in two populations with high or low cell density to a control of low cell density where exogenous CSP was added to artificially synchronise competence development. Samples were taken every 10 min to measure growth (OD_492_), competence development (*P_ssbB_*::*luc*) and the ability to transform during a 10 min or 20 min time window, with DNase I used to prevent subsequent DNA internalisation and resulting transformation after these incubation times (Figure S6A). In the artificially synchronised control, CSP was added at the X_A_ time (70 minutes), identified using P*_ssbB_*::*luc* (Figure S6B). This control leads to an immediate coordination of competence, with a 5-log jump in transformants, mimicking classical QS (Figure S6C). The percentage of transformants was compared for each condition at each time point throughout growth, showing bimodality of the population and correlation of the exponential rate of competence development with cell density (Figure S6C). To further highlight the exponential propagation of competence development, we calculated the difference in transformants obtained at each time point between 20 min and 10 min incubation with tDNA (Δtransformants_(20-10)_) ^74^, since any increase in transformants 20 min after CSP addition compared to 10 min after proves the propagation of competence through a population and disproves a synchronous induction. A sequence of exponentially increasing Δtransformants_(20-10)_ values was observed in both high and low density inocula, with a rate higher in the high density inocula (Figure S6D). In addition, these transformation assays reveal transformant cells prior to competence propagation, which may represent self-induction of a sub-population of cells, highlighting a bimodal mode of development.

#### Mitomycin C creates a self-induced competent cell fraction

The competence stress induction experiment was repeated with MMC rather than streptomycin (Figure 2). It is of note that induction of competence by MMC at 60 ng mL^-1^ leads to a loss of correlation between cfu and OD measurement (Figure S7A, right panels). MMC, via its crosslinking activity, affects chromosome organisation and modifies cell shape ^75^. Observation by microscopy 90 min after antibiotic addition showed that contrary to streptomycin, MMC promotes chaining of 70% of the cells and kills nearly 30% of cells (Figure S7BCD). In these conditions, MMC leads to the disappearance of the self-induced cell fraction that is observed between 80 and 160 min without antibiotic addition (Figure S8B). On the other hand, our results suggest that MMC-mediated stress generates self-inducing cells in the surviving population, leading to competence propagation even if some self-induced cells may not survive.

## Supplementary figure legends

**Figure S1:**
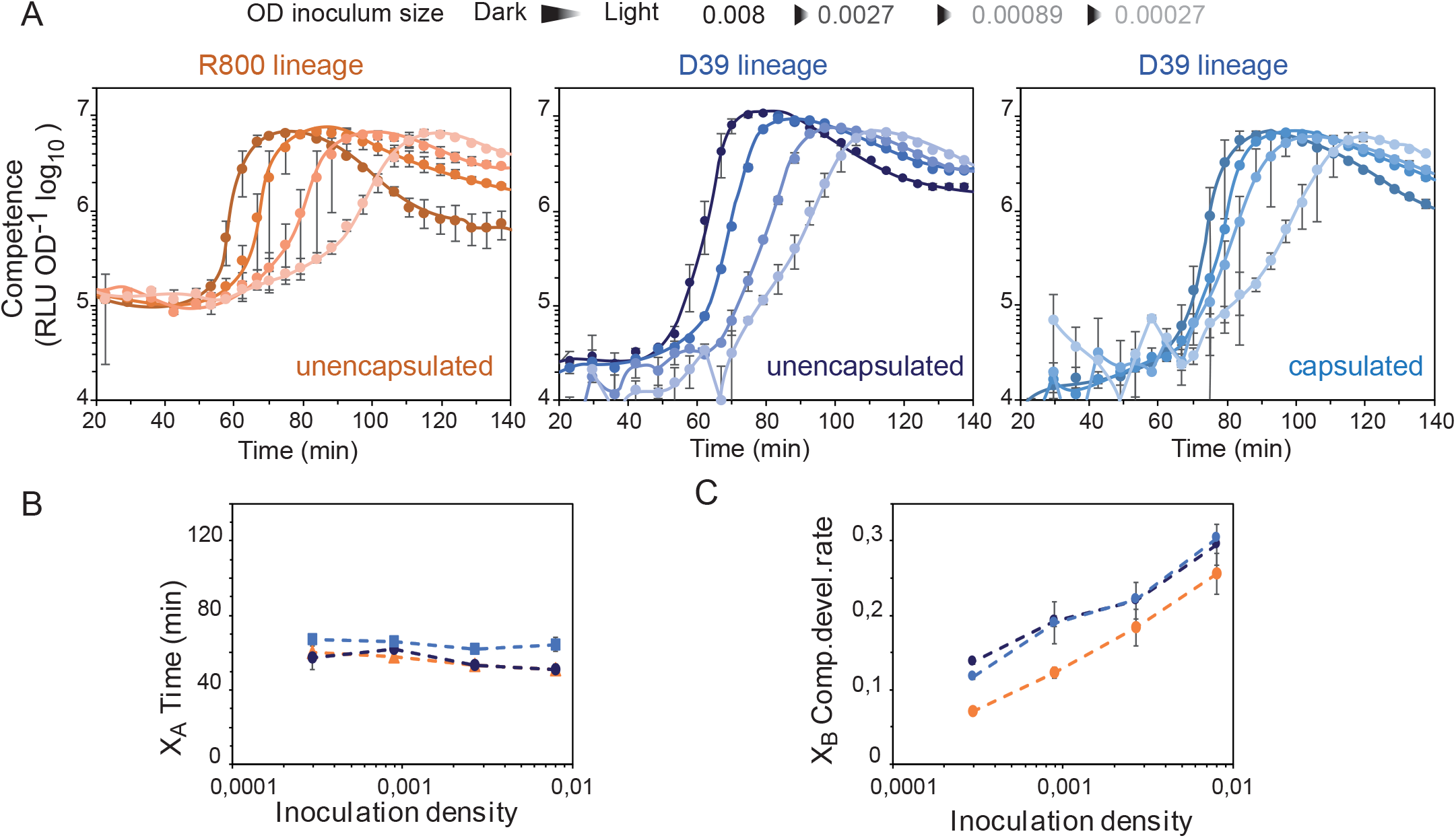
Competence proceeds by propagation irrespective of capsule presence. (A) The P*_ssbB_*::*luc* fusion was used to monitor competence development throughout growth in the unencapsulated R800 lineage (orange) and capsulated and unencapsulated variants of the virulent D39 lineage (light blue and dark blue respectively). Pre-culture stocks at OD_550_ 0,4 were inoculated at 50-, 150-, 450- and 1350-fold dilutions giving inocula ranging from OD_550_ 0,008 (dark) to 0,00027 (light). Specific RLU OD^-1^ readings are reported for each strain at each inoculum. (B) Mean X_A_ times for different densities of inoculum of strains remain constant. (C) Mean competence development rate (X_B_) for different densities of inoculum of strains correlates to cell density. Standard deviation calculated from triplicate repeats.

**Figure S2:**
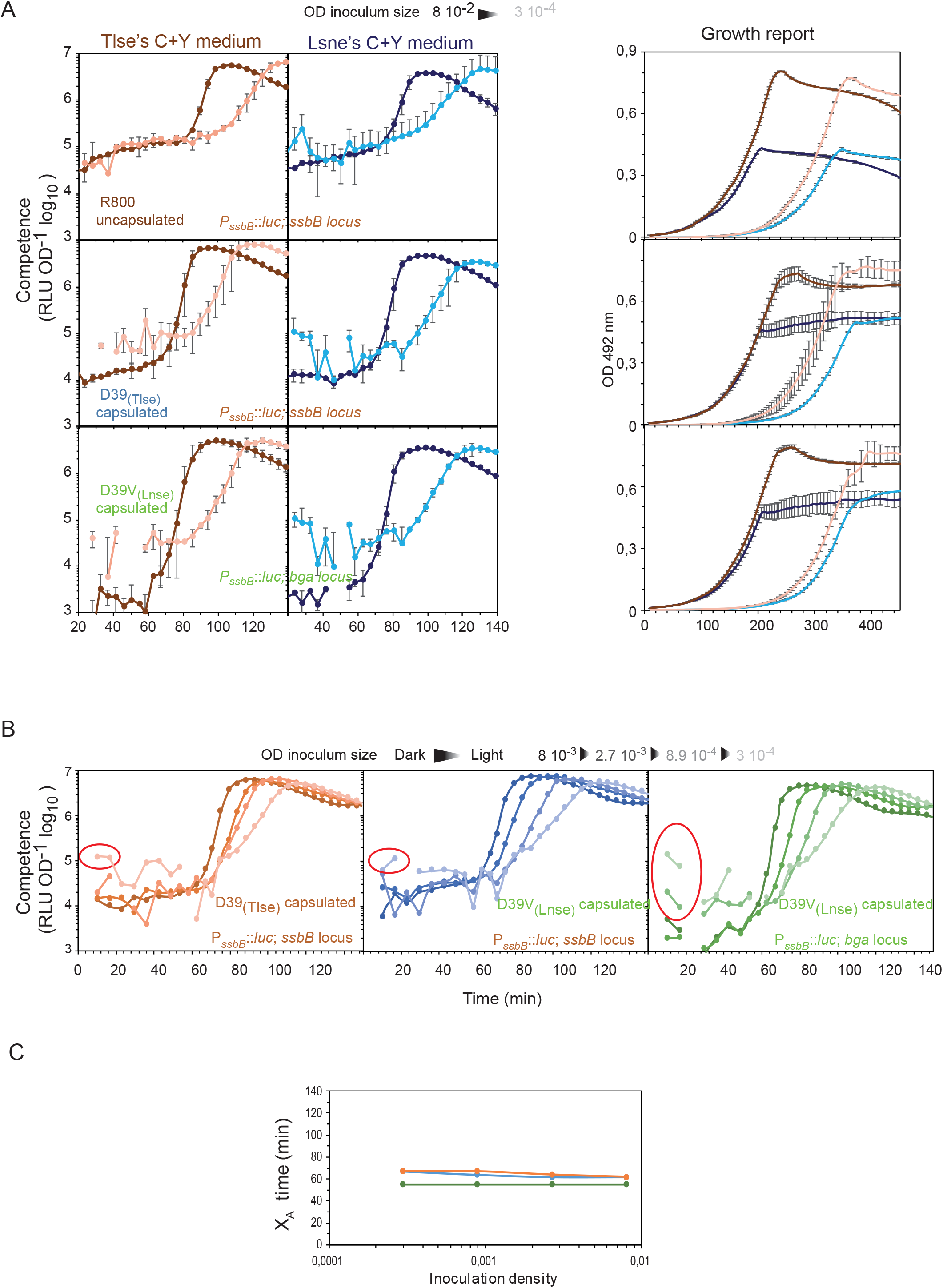
D39 strains or medium used do not alter the propagative behaviour of pneumococcal competence. (A) Comparison of competence development (RLU OD^-1^) of three strains in different C+Y medium prepared in Toulouse (Tlse C+Y medium) (brown scaled colours) and Lausanne (Lsne C+Y medium) (blue scaled colours). Strains used: two from the Toulouse collection (Tlse), the unencapsulated R800 lineage (R895, brown colour) and D39 (TD277, blue colour) both with P*_ssbB_*::*luc* at the *ssbB* locus but also ssbB^+^ at the *ssbB* locus, and the third one from Lausanne (Lsne), D39V with P*_ssbB_*::*luc* in the *bgaA* locus, Δ*bgaA* (DLA3) (green colour). Left panels show competence development in specific activity (RLU OD^-1^) and right panels show growth (OD_492_). Standard deviations are indicated for all experiments based on triplicate repeats. (B) P*_ssbB_*::*luc* in the *bgaA* locus is a poor reporter of spontaneous competence propagation compared to P*_ssbB_*::*luc* at the *ssbB* locus. D39_(Tlse)_ containing P*_ssbB_*::*luc* at the *ssbB* locus is compared to D39V(Lsne) containing P*_ssbB_*::*luc* at the *ssbB* or *bgaA* loci. Red circles point out data that are out of range of the sensitivity of the Thermofisher Varioskan Flash luminometer (see also Figure S3). (C) Deduced X_A_ time of each strain for each inoculum size in panel B. Individual data shown representative of triplicate repeats showing similar results.

**Figure S3:**
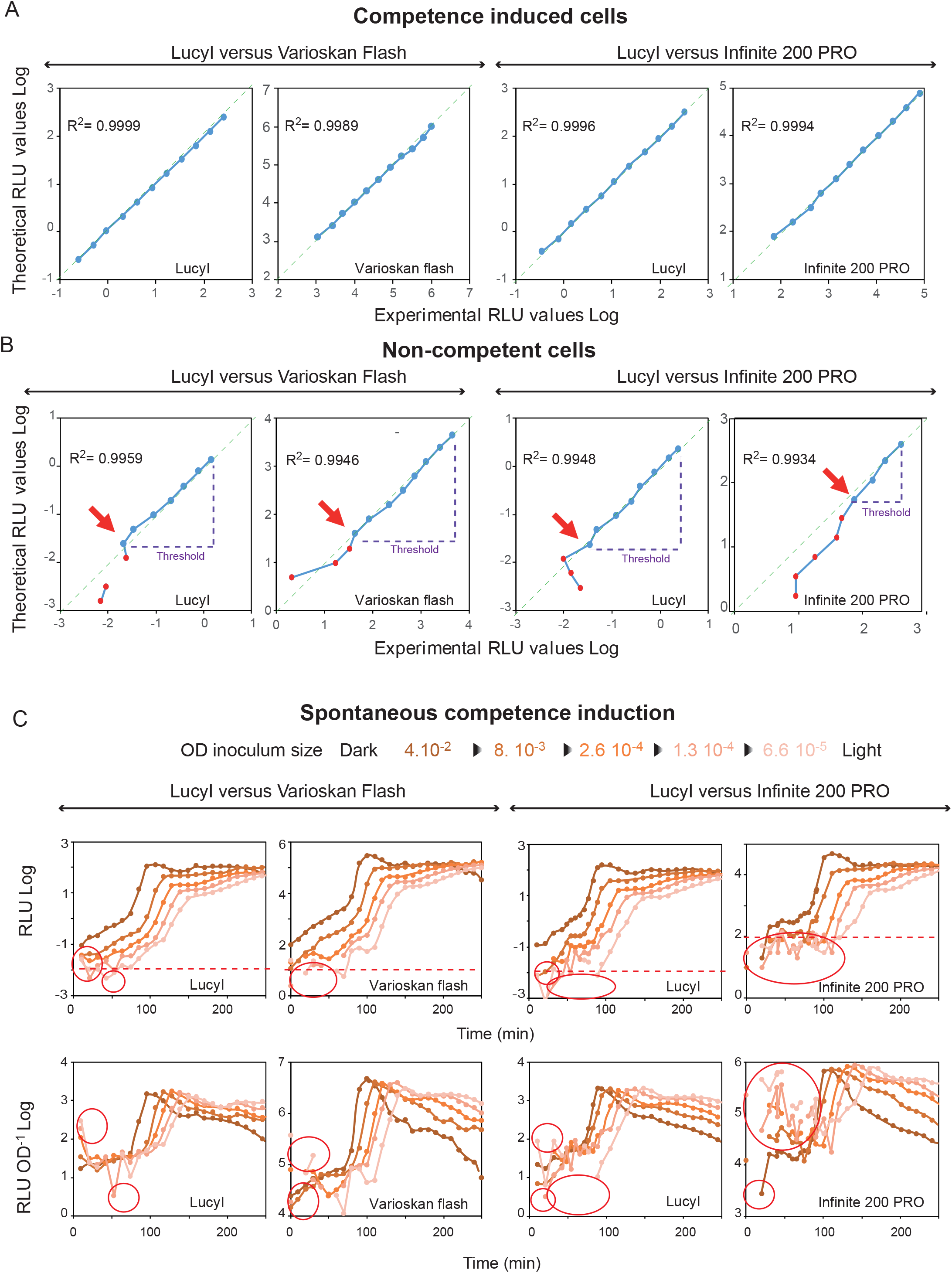
Comparing the RLU sensitivity threshold of luminometers. Comparing sensitivity of luminometers detecting competent (A) and non-competent cells (B) in serially diluted samples. The luminometers used in each study (Thermo Varioskan Flash ^39^, Tecan Infinite 200 PRO ^38^) were individually compared to a reference luminometer (LucyI, Anthos). The graphs represent the experimental RLU value plotted (x axis) against the theoretical calculated value (y axis) obtained by dividing the highest measured value by two-fold successively. The RLU background is obtained by measurement of an equivalent volume of sterile medium. The green dotted diagonal allows visualisation of whether the luminometer is accurate or not. Red dots represent values considered out of range. The threshold limit of each luminometer is indicated by the dotted purple lines. (C) Comparison between luminometers to detect the first wave of spontaneous competence development. The data out of range of the instrument are circled in red. The dotted horizontal red line represents the approximative threshold limit of the luminometer. We were unable to calculate the X_A_ for almost all cell densities with the Tecan Infinite 200 PRO luminometer.

**Figure S4:**
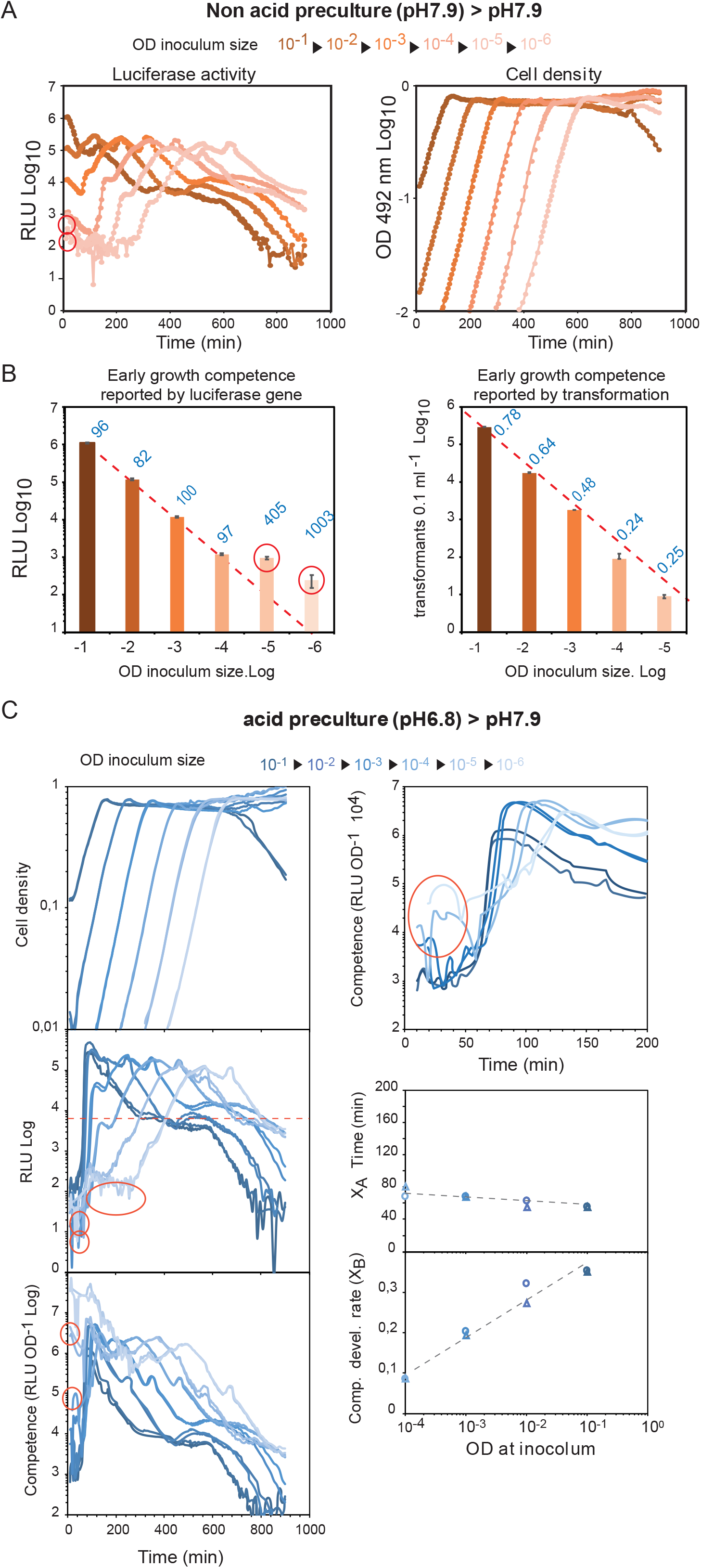
Dilution of competent cells does not abolish competence. We repeated the experiment described in Figure 2B of the study which suggested that dilution of competent cells abrogated competence ^38^. (A) RLU (left) and OD_492_ (right). Red circles, data out of range of the instrument at the beginning of the experiment. (B) Left graph, mean RLU between 10 and 14 minutes after inoculation. The blue number on each column corresponds to calculated RLU OD^-1^ that is independent of the cell dilution. Right graph, total numbers of streptomycin resistant transformants in cfu per 0.1 mL for each inoculum size at 10 min of the growth culture (see colour guide in panel A). The blue number on each column reports the percent of transformants for each inoculum size. Red dotted lines represent the theoretical expected value deduced from the highest inoculum size if dilution does not affect competence. Standard deviations are represented. (C) We repeated the experiment described in Figure 2A of the same study ^38^. DLA3 displays propagative competence induction. Left graphs: upper panel, cell density growth curves (OD_492_); middle panel, RLU; bottom panel, specific activity (red circles show data out of range of the instrument). The horizontal dotted red line in the middle panel corresponds approximately to the RLU competence threshold previously described ^38^. Right graphs: top panel, zoom of first 200 min of bottom left graph (red circle shows data out of instrument range); bottom panel, deduced X_A_ time and the deduced competence development rate (X_B_) of each strain for each inoculum size. Triangles and circles represent two different assays per inoculum size. The dotted grey line corresponds to the tendency between the assays. The correlation coefficient R^2^ for each exponential regression calculation (X_B_) was done on a minimum of five consecutive values and found to be between 0.91 and 0.99. All the experiments described above were reproduced with the TD288 strain and gave similar results.

**Figure S5:**
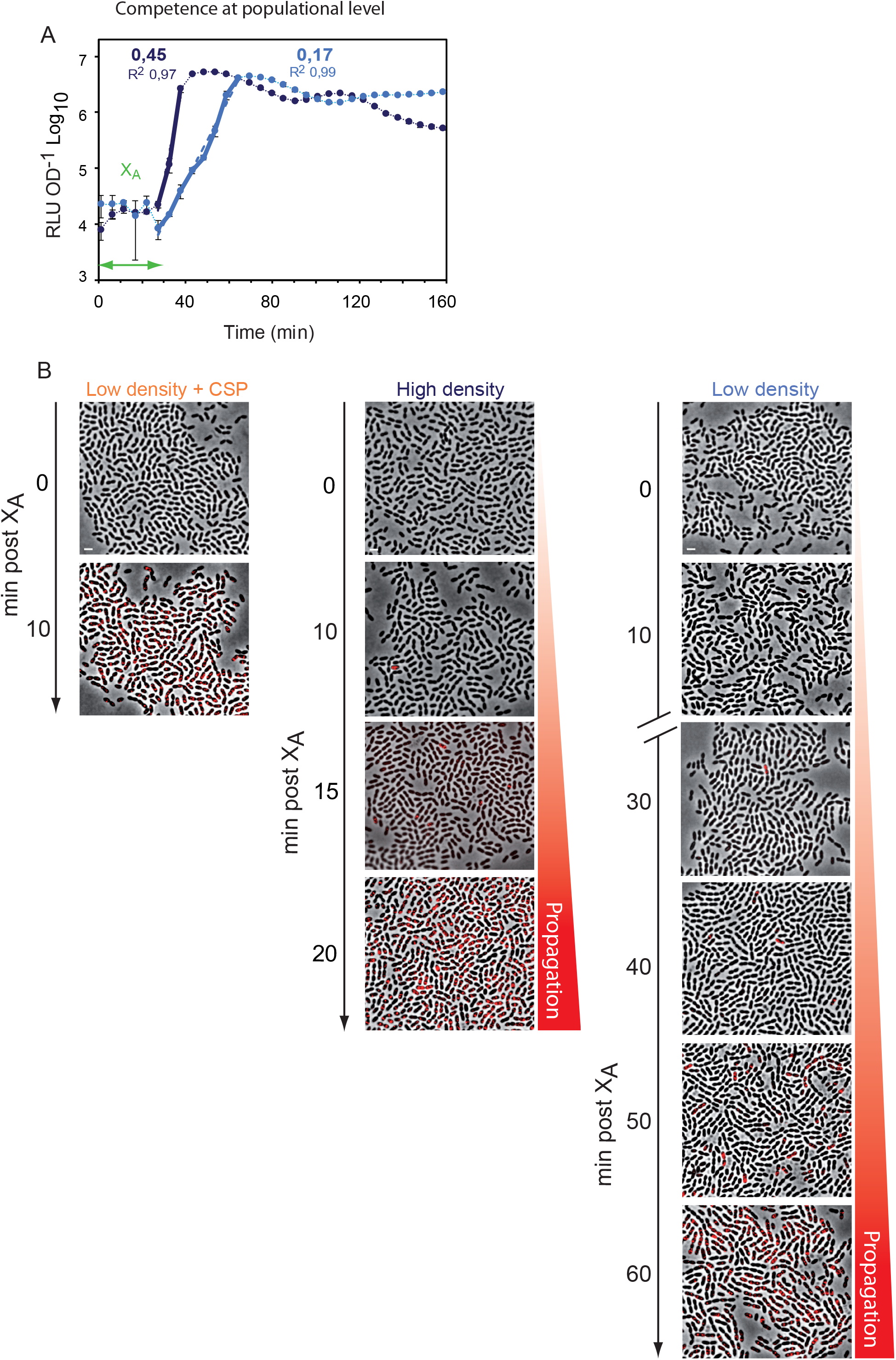
Competence propagation tracked by microscopy. Tracking binding of fluorescent tDNA to competent cells in different densities of inocula supports the propagation model. (A) Calculation of X_A_ time by tracking P*_ssbB_*::*luc*, to determine when to begin observation of spontaneous competence development by microscopy. The graph represents the specific activity (RLU OD^-1^) readings every 10 minutes. The X_A_ time (double-headed green arrow) is reported. The competence development rate values (X_B_) are calculated with data restricted to the thick lines with the R^2^ correlation coefficient. X_A_ time determined as 30 min in these conditions. (B) Fluorescent images representative of cells incubated with 285-bp Cy3-DNA during the time course as in Figure 1E, white scale bar 2 µm. Individual data shown representative of triplicate repeats showing similar results.

**Figure S6:**
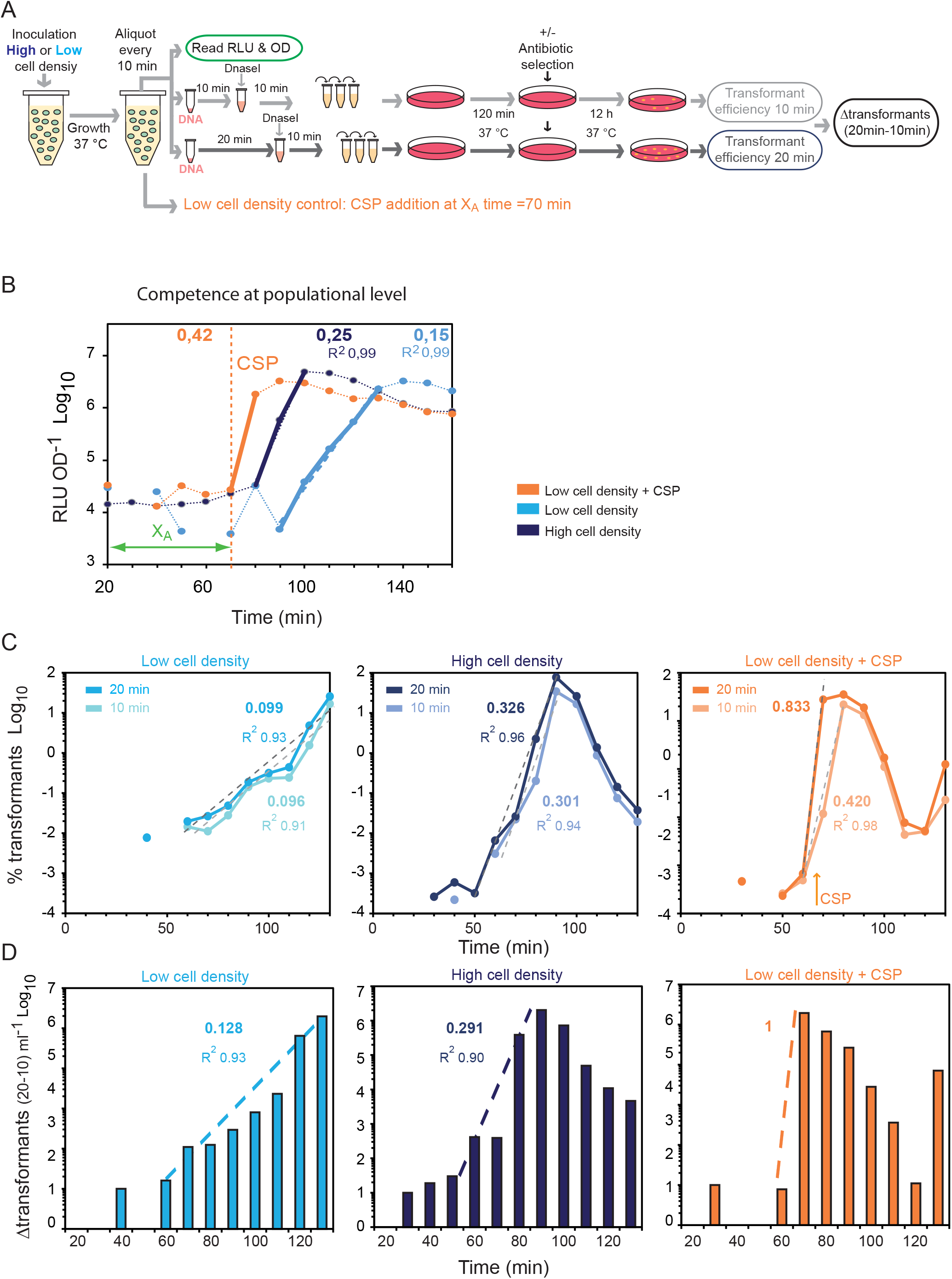
Self-Induction and propagation (SI&P) observed at single cell level. (A) Schematic of transformation assays to follow competence development. The difference in transformants between 20 min exposure and 10 min exposure to transforming DNA (Δtransformants_(20 min-10 min)_) reports the amount of cells shifting to competence during the period between 10 min and 20 min after the sampling. (B) Competence propagation observed via P*_ssbB_*::*luc* as control in parallel with transformation assay using TD277. This strain was pre-cultured in non-permissive medium and was used to generate a low cell density (2.9 10^-4^ OD, light blue curves) and a high cell density (8 10^-3^ OD, dark blue curves) inoculum size in permissive medium. The orange curves correspond to the experiment with addition of CSP (100 ng mL^-1^) to a low density culture after 70 minutes of growth (orange small dotted vertical line). The graph represents the specific activity (RLU OD^-1^) readings every 10 minutes of the experiment. The competence development rates with associated X_B_ values calculated from the exponentials (thick curves) are given with the R^2^ correlation coefficient (note that for the assay with CSP only two points can be taken into account). (C) Transformation efficiency of TD277 reported during growth of cells in permissive medium. Left panel, low cell density inoculation with 10 min (light cyan) or 20 min (cyan) exposure to tDNA; middle panel, high cell density inoculation with 10 min (light blue) or 20 min (dark blue) exposure to tDNA; right panel; low cell density inoculation with addition of 100 ng mL^-1^ CSP after 70 min of growth, with 10 min (light orange) or 20 min (orange) exposure to tDNA. Rate of transformant increase with correlation coefficient (R^2^) is reported for each condition. Note that the first transformants observed prior to propagation may represent self-induced competent cells. Colour scheme as in panel B. (D) Comparing difference between transformant levels at 10 and 20 minutes post-CSP at different cell densities demonstrates exponential propagation as the mode of competence development. The Δtransformants_(20 min-10 min)_ is reported for each sampling for each assay. The rate of increase of Δtransformants_(20 min-10 min)_ is reported with the R^2^. Dashed lines show data range used for rate calculations. Colour scheme as in panel B. Data shown representative of three individual repeats.

**Figure S7:**
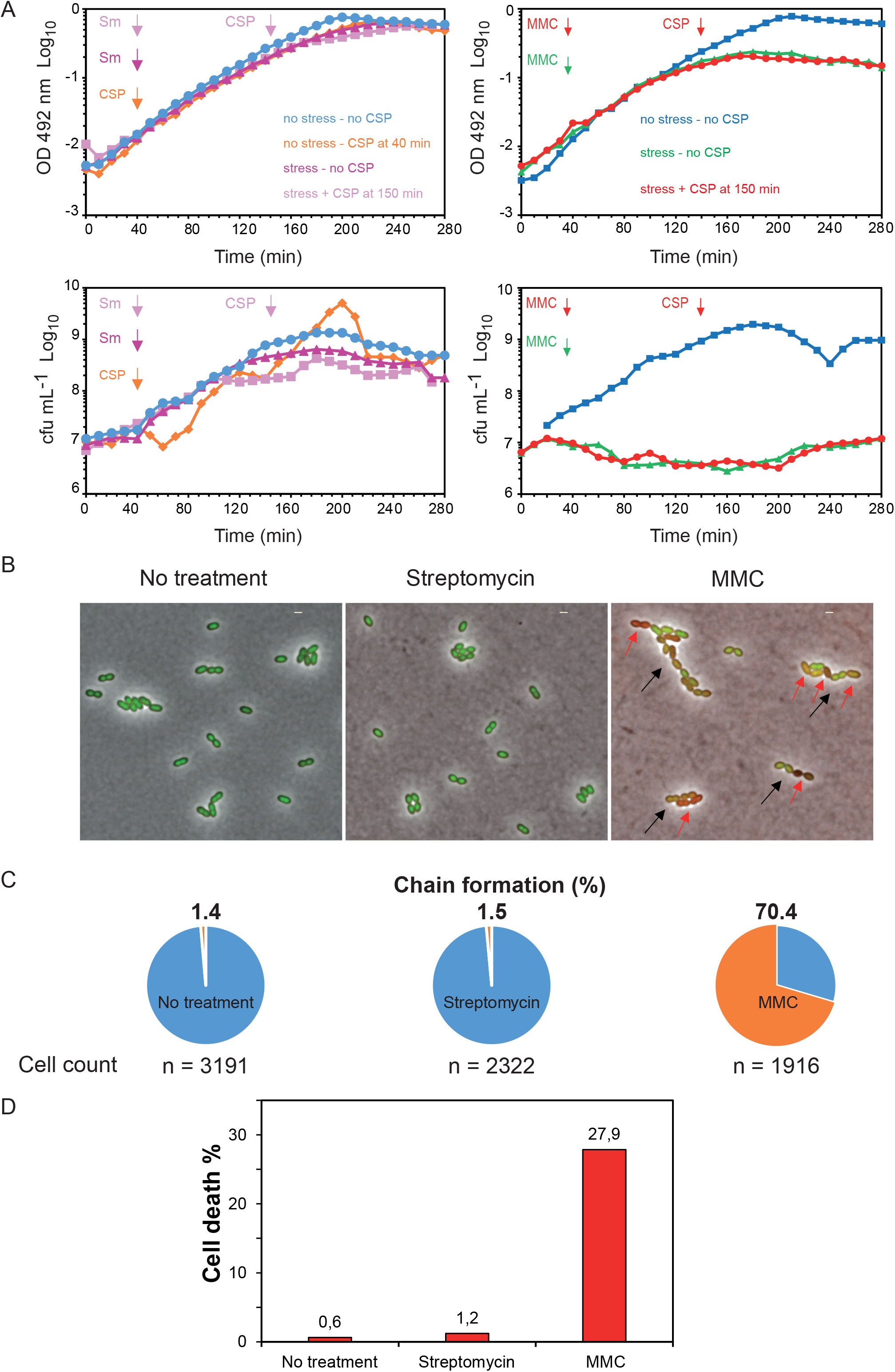
Mitomycin C induces cell chaining and cell death. (A) Growth and viability measurements in experiments of competence induction by antibiotics. Top panels, OD followed during experiments. Bottom panels, cfu mL^-1^ obtained during the experiments presented in Figure 2 with streptomycin in Figure S8 with MMC. (B) Visualisation of cell chaining and survival by microscopy using Live/Dead assay. Black arrows, cell chaining, red arrows, cell death. (C) Percentage of cell chaining detected for each condition. (D) Percentage of cell death observed 90 minutes after time of antibiotic addition.

**Figure S8:**
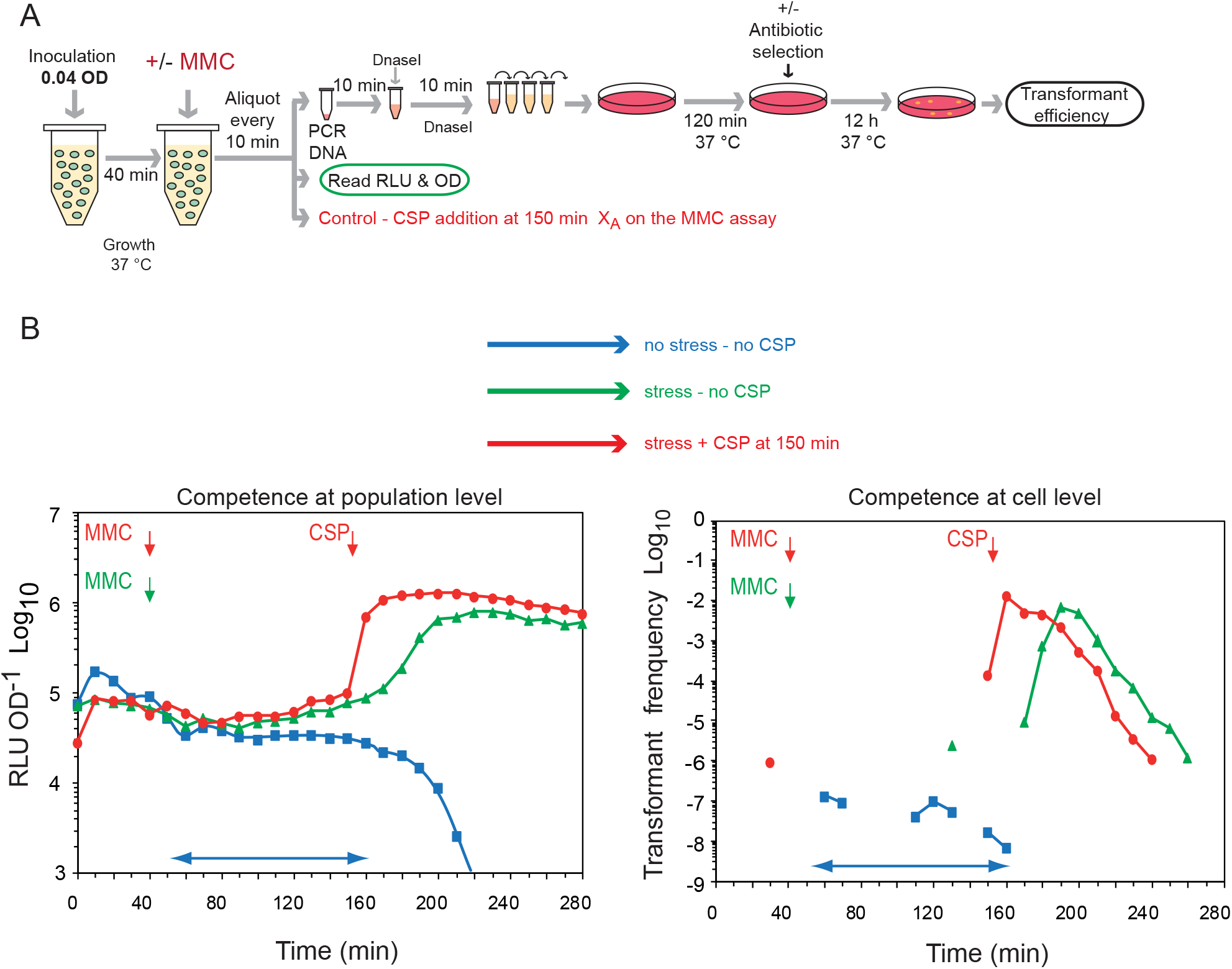
Competence propagation induced by MMC. (A) Experiment conducted as described in Figure 2, but with MMC replacing streptomycin. Schematic representation of experiment carried out to explore whether competence induction by exposure to sub-lethal concentrations of MMC follows a SI&P mode of transmission. (B) MMC at 60 ng mL^-1^ was added after 40 minutes of growth (red and green curves). CSP was added after MMC at 150 minutes (red curves) of growth at the concentration of 100 ng mL^-1^. The blue curve corresponds to the control without any addition. The left graph reports the populational competence tracked by RLU OD^-1^ and the right graph reports individual kanamycin resistant transformant tracking. Arrows represent time of addition of CSP or antibiotic. Individual data shown representative of triplicate repeats showing similar results. The blue arrowed lines highlight a period with detection of low transformant levels (right graph) revealing a self-induced cell fraction, without detection of populational competence propagation (left graph).

**Figure S9:**
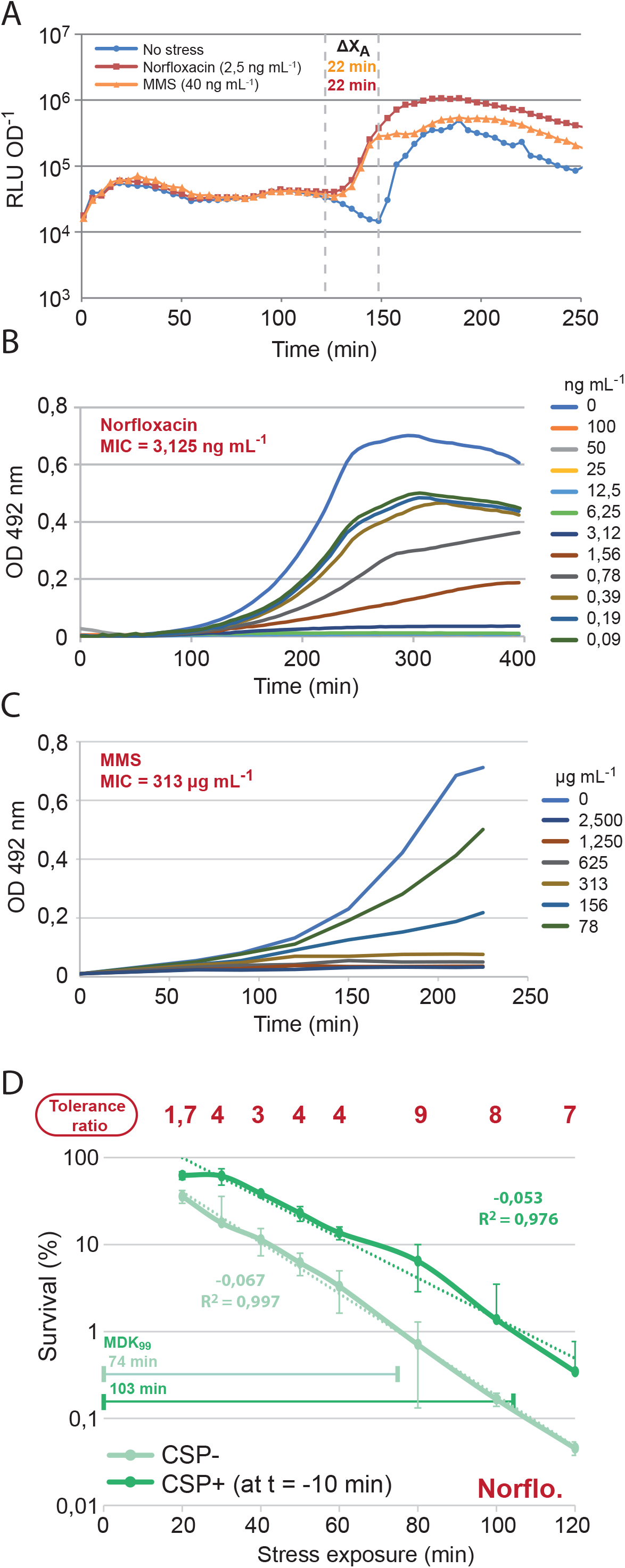
MIC calculation, competence induction and survival time course of Norflo and MMS. (A) Exposure to Norflo or MMS at sub-MIC levels reduces the X_A_ period of competence in R895 *comC^+^*cells able to spontaneously develop competence. Competence was visualised using P*_ssbB_*::*luc*. Stresses added at 0 min. Data representative of triplicate repeats. (B) Growth of pneumococci in a gradient of norfloxacin concentrations to determine MIC, defined as the lowest concentration blocking pneumococcal growth. Data representative of triplicate repeats. (C) Growth of pneumococci in a gradient of MMS concentrations to determine MIC, defined as in panel B. Data representative of triplicate repeats. (D) Time-course of survival of competent (dark green) and non-competent (light green) R1501 cells exposed to norfloxacin. Tolerance ratios calculated as in Figure 3A. Dotted lines represent exponential fits with exponential rates and R^2^ values provided. MDK_99_ values represent time taken to kill 99 % of the population. Means and standard deviations calculated from triplicate repeats.

**Figure S10:**
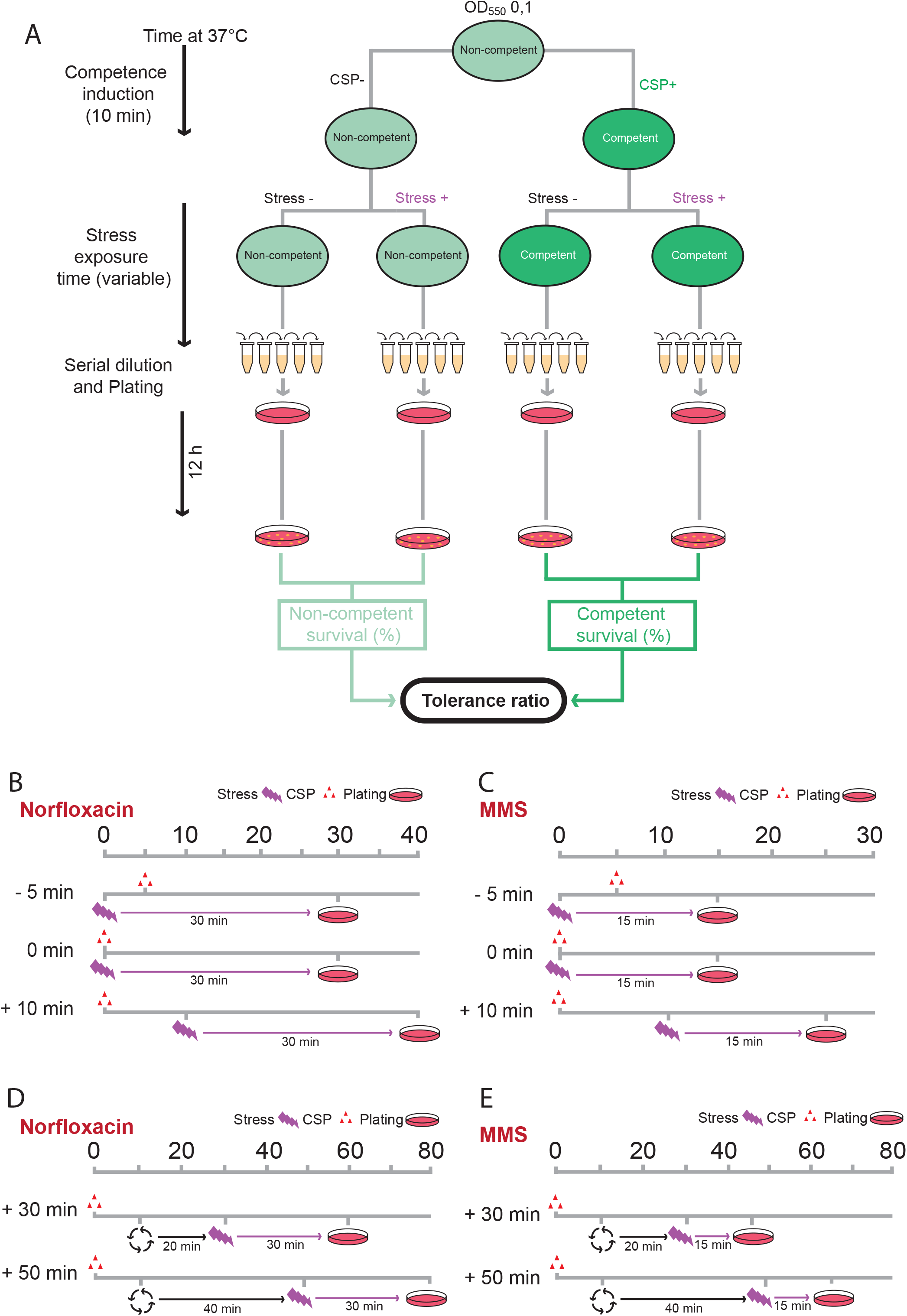
Schematic representations of survival assays. (A) Pre-competent cells were grown to OD_550_ 0,1 and split into two cultures, with one half induced to competence by addition of 100 ng mL^-1^ synthetic CSP. After 10 min at 37 °C, cultures were again split and exogenous stress was added or not, before further incubation at 37 °C for variable time. Cells were then serially diluted and plated before a final incubation at 37 °C for 12 h. Comparing cfu from stressed and non-stressed conditions allowed calculation of survival ratios for competent and non-competent cells, and comparison of these ratios produced the tolerance ratio, revealing the effect of competence on tolerance to a particular stress. (B) Schematic representation of timings of Norflo survival assays exploring exposure to stress at different times relative to CSP addition. (C) Schematic representation of timings of MMS survival assays exploring exposure to stress at different times relative to CSP addition. (D) Schematic representation of timings of Norflo survival assays exploring exposure to stress at time points after CSP addition. (E) Schematic representation of timings of MMS survival assays exploring exposure to stress at time points after CSP addition.

**Figure S11:**
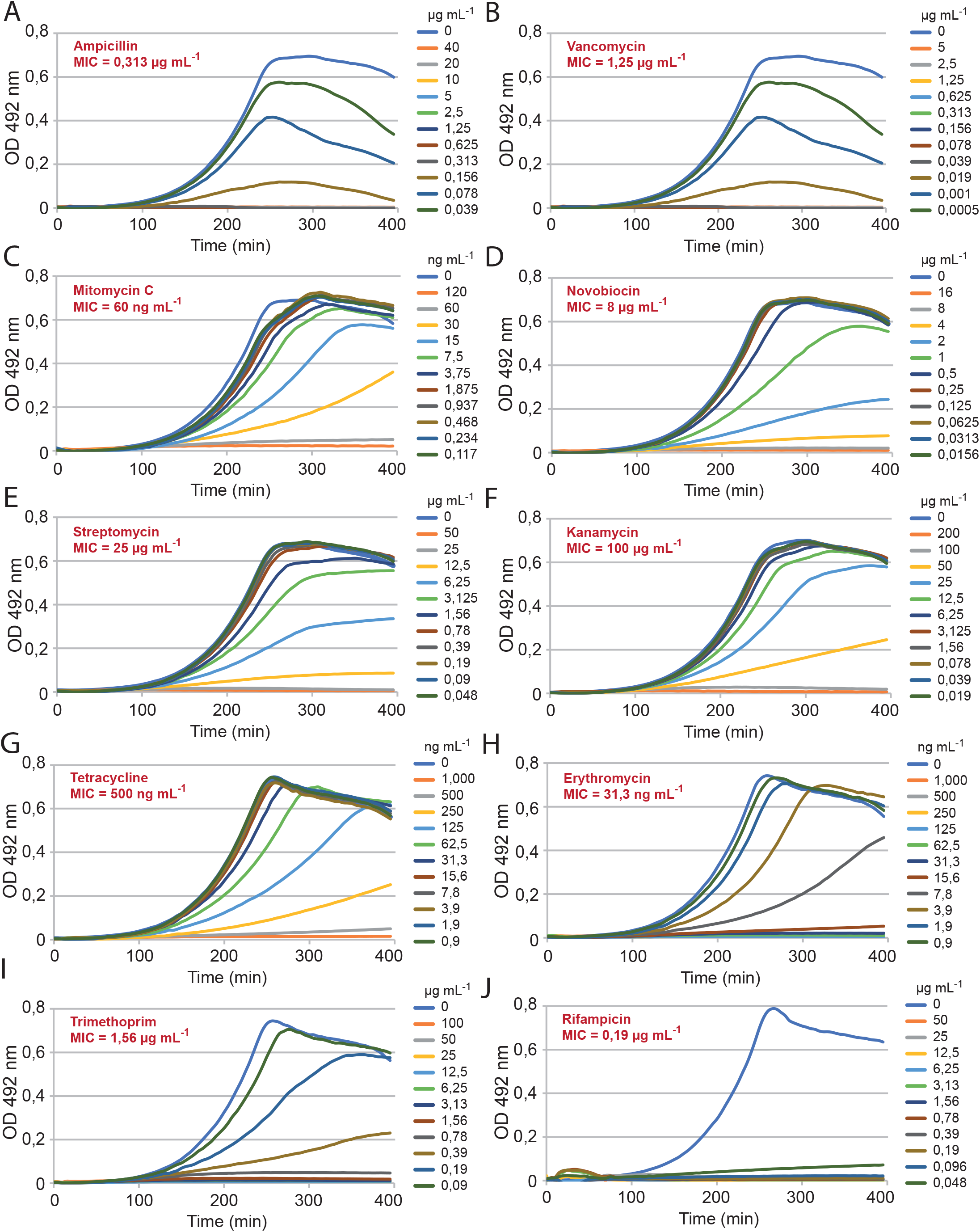
Growth curves for calculation of MIC of tested stresses. Pneumococci in a gradient of stress concentrations to determine MIC, defined as in Figure S9. Data representative of triplicate repeats. (A) Ampicillin. (B) Vancomycin. (C) MMC. (D) Novobiocin. (E) Streptomycin. (F) Kanamycin. (G) Tetracycline. (H) Erythromycin. (I) Trimethoprim. (J) Rifampicin.

**Figure S12:**
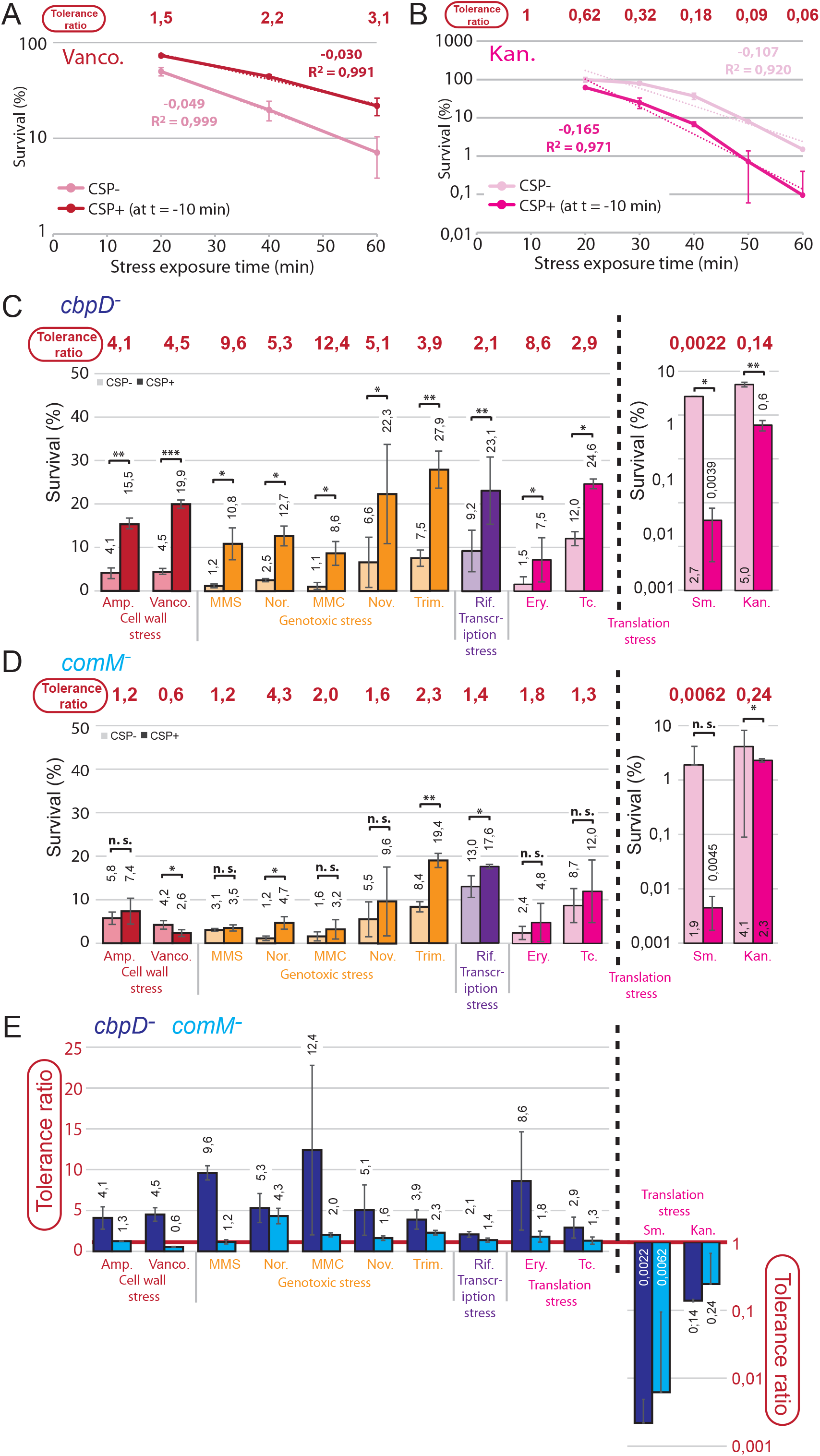
Survival assay control experiments in wildtype, *cbpD^-^* and *comM^-^* cells. (A) Time-course of survival of competent (dark red) and non-competent (light red) R1501 cells exposed to vancomycin. Tolerance ratios calculated as in Figure 3A. Dotted lines represent exponential fits with exponential rates and R^2^ values provided. (B) Time-course of survival of competent (dark pink) and non-competent (light pink) R1501 cells exposed to kanamycin. Tolerance ratios calculated as in Figure 3A. Dotted lines represent exponential fits with exponential rates and R^2^ values provided. (C) Survival of competent (dark) and non-competent (light) *cbpD^-^*cells (R4951) exposed to various stresses for 60 min starting at +10 min relative to CSP addition. Experimental procedures and representations as in Figure 4A. *, p < 0,05; **, p < 0,01; ***, p < 0,005. (D) Survival of competent (dark) and non-competent (light) *comM^-^* cells (R4950) exposed to various stresses for 60 min starting at +10 min relative to CSP addition. Experimental procedures and representations as in Figure 4A. n.s., non-significant, p > 0,05; *, p < 0,05. (E) Comparison of tolerance ratios of *cbpD^-^* and *comM^-^* cells, with values as in Figure 4C.

**Figure S13:**
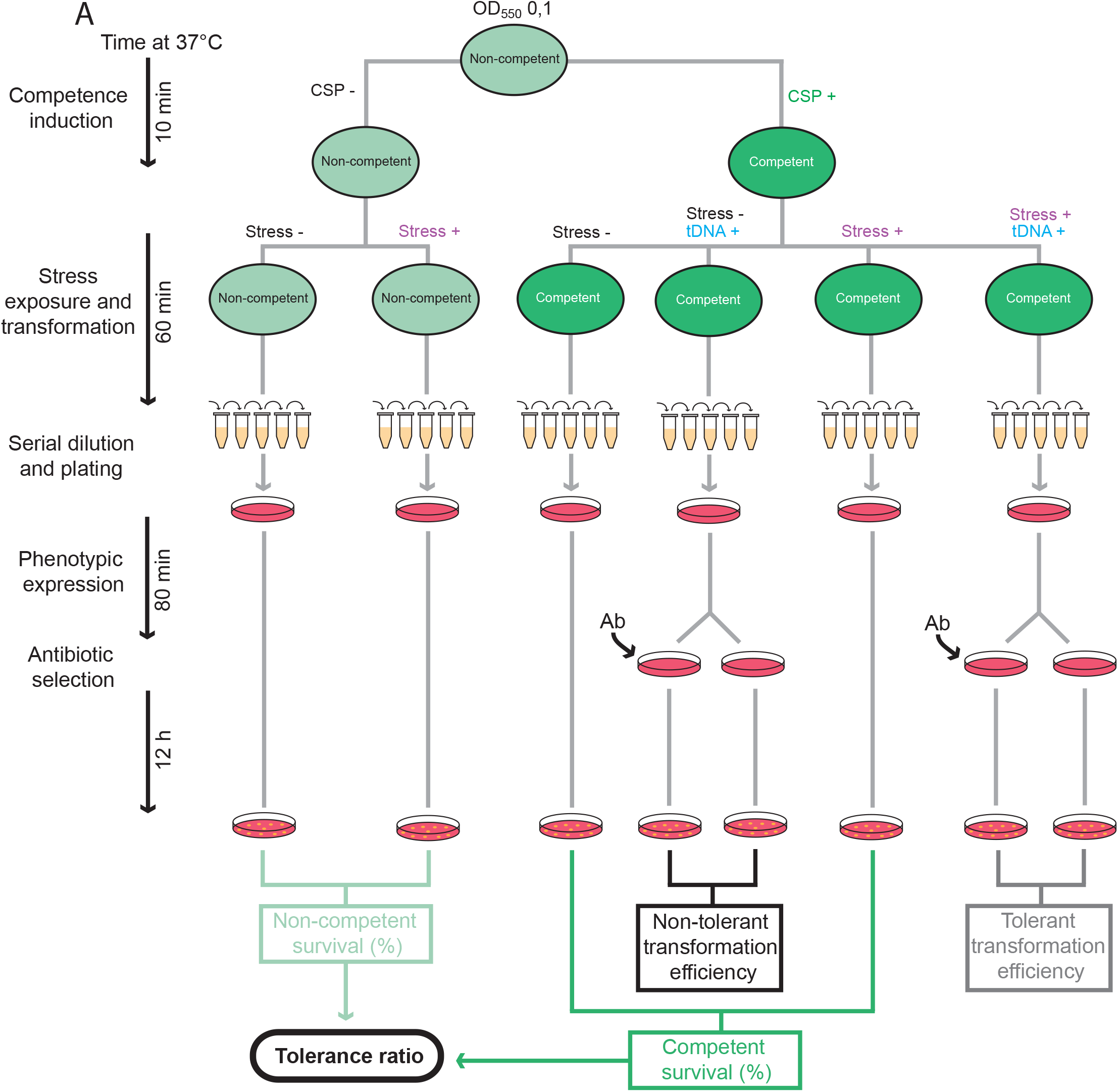
Schematic representation of transformation assay. (A) Pre-competent R3369 cells were grown to OD_550_ 0,1 and split into two cultures, with one half induced to competence by addition of synthetic CSP. After 10 min at 37°C, cultures were again split and tDNA and exogenous stress were added as shown, before a further 60 min incubation at 37°C. Cells were then serially diluted and plated before 80 min of phenotypic expression at 37°C. A second layer of medium with selective antibiotic was added to desired tDNA^+^ plates to select for transformants. Comparison of cfu in stress +/- conditions in CSP+ or CSP- cells determined the tolerance of competent and non-competent cells in the face of the stress. Comparing these values revealed the effect of competence on stress tolerance via the tolerance ratio. Transformation efficiencies of stress +/- populations were determined by comparing cfu on selective and non-selective plates.

